# Myeloid HDAC7 drives liver inflammation and systemic glucose dysregulation during diet-induced obesity

**DOI:** 10.64898/2025.12.04.691405

**Authors:** Yizhuo Wang, Divya Ramnath, Kaustav Das Gupta, P. Prakrithi, Kavita Bisht, Gregory C. Miller, Zherui Xiong, Yujun Wan, Ellen N. Tejo, James E.B. Curson, Rishika Abrol, Sahar Keshvari, Kimberley S. Gunther, Jordan D. Atkinson, Zhixuan Loh, Jessica A. Engel, Christian R. Engwerda, Sabrina Sofia Burgener, Kate Schroder, David P. Fairlie, Andrew D. Clouston, Elizabeth E. Powell, Katharine M. Irvine, Mitchell A. Sullivan, Jean-Pierre Lévesque, Quan Nguyen, Matthew J. Sweet, Denuja Karunakaran

## Abstract

**Objectives:** Histone deacetylase 7 (HDAC7), a classical HDAC family member, promotes LPS-inducible glycolysis and inflammatory mediator production in macrophages, innate immune cells that contribute to pathology in metabolic diseases. Here, we investigated myeloid HDAC7 functions in obesity-driven metabolic disease.

**Methods:** We used gain- and loss-of-function genetic approaches in mice to investigate myeloid HDAC7 functions in hepatic inflammation and metabolic disease, as well as associations with hepatic gene signatures characteristic of advanced chronic liver disease (CLD).

**Results:** Transgenic expression of *Hdac7* in myeloid cells increased liver inflammation and liver mRNA levels of *Ccl2* and *Il1b*, key inflammatory mediators linked to CLD. Liver glycogen levels were also decreased, another feature of CLD. Transgenic expression of *Hdac7* in myeloid cells mimicked the hepatic inflammatory phenotype that was observed in mice fed a high fat, high cholesterol, and high sucrose (HFHCHS) diet, an obesity model that mimics some features of metabolic dysfunction-associated steatotic liver disease. In myeloid *Hdac7* transgenic mice fed a HFHCHS diet, relative weight gain was increased, fasted glucose levels were elevated and glucose tolerance was dysregulated by comparison to control mice. Conversely, fasted blood glucose levels were reduced and glucose tolerance was improved in myeloid *Hdac7*-deleted mice on a HFHCHS diet. *HDAC7* mRNA levels were also elevated in the livers of people with advanced CLD and spatial transcriptomics revealed that myeloid HDAC7 directs hepatic gene signatures characteristic of advanced CLD.

**Conclusions:** Myeloid HDAC7 contributes to hepatic inflammation and systemic glucose dysregulation in a mouse model of obesity and liver inflammation.

## Introduction

Metabolic dysfunction-associated steatotic liver disease (MASLD) is a common cause of chronic liver disease (CLD) that is associated with substantial morbidity and mortality^1^. MASLD progressing to metabolic dysfunction-associated steatohepatitis (MASH) is the major cause of cirrhosis, especially in developed countries^2^. MASLD represents a spectrum of conditions, ranging from fatty liver (steatosis) without significant inflammation to MASH, where liver injury, inflammation, and/or fibrosis accompany steatosis^3^. The thyroid hormone receptor-β agonist Resmetirom^4^ and the glucagon-like peptide-1 (GLP-1) receptor agonist semaglutide^5^ are approved in the USA as therapies for patients with moderate to advanced liver fibrosis. Other therapies are currently under clinical evaluation^6^, including tirzepatide, a dual agonist for GLP-1 and glucose-dependent insulinotropic polypeptide receptors^7^. However, these therapies are likely to be effective in only a sub-set of MASH patients. The development of more effective treatments requires a better understanding of the complex underlying mechanisms that drive MASH and the employment of approaches that target multiple inflammatory and fibrotic pathways.

Macrophages are key innate immune cells that sense and respond to endogenous and exogenous danger signals, enabling them to detect perturbations in homeostasis^8^. These cells are extremely heterogeneous, exhibiting diverse phenotypes depending on the specific environmental conditions encountered^9^. Liver macrophages, comprising both tissue-resident Kupffer cells and monocyte-derived macrophages, play important roles in liver homeostasis and in responding to environmental signals. Kupffer cells clear endotoxin and phagocytose bacterial pathogens, dying cells and other debris in the hepatic parenchyma^10, 11^, and also release inflammatory cytokines upon activation via the pattern recognition receptors, such as toll-like receptor (TLR) 4^12^. Recruited monocytes can also differentiate into macrophages in the liver, either promoting wound healing and tissue repair when the injury-inducing agents have been removed^13, 14^ or perpetuating inflammation and pathology during active disease^15, 16^. A failure to resolve macrophage activation leads to dysregulated production of inflammatory mediators that contribute to pathological conditions, including liver fibrosis^10^. For example, elevated TNF levels in inflamed livers contribute to fibrogenesis^17^, the chemokine CCL2 promotes lipid accumulation in hepatocytes leading to obesity-induced liver steatosis^18^ and MASH progression^19^, and inflammasome-mediated IL-1β production is associated with collagen deposition in MASH patients^20^. Targeting of such inflammatory mediators can attenuate progression of liver disease in mouse models. For example, inhibition of IL-1 receptor signalling^21^ or deletion of *Il1b* that encodes IL-1β^22^ in mouse models limited MASH progression.

To fuel proinflammatory responses, aerobic glycolysis is rapidly increased in TLR-activated macrophages^23, 24^. This is linked with TCA cycle reprogramming, ultimately resulting in the generation of acetyl-CoA that enables histone modifications for inducible expression of inflammatory genes^25, 26^. Signalling molecules that connect glycolysis and inflammation such as the glycolytic enzyme pyruvate kinase isoform 2 (PKM2) have been implicated in the pathogenesis of MASH. For example, elevated levels of PKM2 in both serum and liver were associated with MASLD severity^27^. PKM2 exists in two forms, a tetramer that is associated with its glycolytic function and a dimeric form that drives inflammatory responses. Tetrameric PKM2 polarises macrophages towards a regulatory or anti-inflammatory phenotype, ameliorating liver inflammation and steatosis^28^. In contrast, nuclear accumulation of dimeric PKM2 in macrophages was reported to promote macrophage polarisation towards a pro-inflammatory phenotype, leading to MASH pathogenesis^29^. Significantly, a group of proteins called the class IIa histone deacetylases (HDACs) promote dimerization and activation of the pro-inflammatory form of PKM2 in macrophages^30^.

HDACs remove an acetyl moiety from the ɛ-amino group of lysine residues of histones, with this generally associated with repression of gene transcription^31^. However, thousands of non-histone proteins are also regulated by lysine acetylation^32, 33^, so HDACs control many cellular processes beyond gene expression. Class IIa HDACs are a subfamily of the classical HDACs that share a conserved C-terminal deacetylase domain and a less conserved N-terminus that interacts with a broad range of cellular proteins^34^, allowing this family to regulate various biological processes in physiological and pathological conditions (reviewed in^35, 36^). One class IIa HDAC with a particularly important function in metabolic and inflammatory processes in macrophages is HDAC7 (reviewed in^37, 38^). In human and murine macrophages, HDAC7 orchestrates LPS-induced glycolysis and the production of key inflammatory mediators IL-1β and CCL2^30^, two inflammatory mediators associated with MASLD/MASH. Interestingly, HDAC7 primarily functions in macrophages responding to submaximal concentration of LPS^39^, suggesting that this class IIa HDAC may be particularly important in driving chronic low-grade inflammation. In contrast, HDAC7 engages the pentose phosphate pathway in macrophages responding to bacterial challenge, with this enabling pathogen destruction and attenuation of inflammation^40^. This highlights a role for HDAC7 in context-dependent danger recognition during inflammation, meaning that predicting its function in different disease settings such as MASLD/MASH is not straightforward. In this study, using myeloid *Hdac7* gain- and loss-of-function mice, we reveal a role for myeloid HDAC7 in liver inflammation and glucose dysregulation in an obesity model that shares some features with MASLD.

## Results

### Myeloid *Hdac7* overexpression promotes liver inflammation in mice

We previously showed that HDAC7 drives production of the pro-inflammatory mediators CCL2 and IL-1β in TLR4-activated macrophages^30, 39^ and that acute LPS-induced systemic inflammation was amplified in MacHDAC7 mice that selectively over-express HDAC7 in myeloid cells^30^. However, whether myeloid HDAC7 drives inflammatory responses and/or pathological changes in the liver has not been examined. To determine if myeloid HDAC7 might contribute to chronic and low-grade liver inflammation, we utilised MacHDAC7 transgenic mice^30^ that overexpress an alternatively spliced, inflammatory form of mouse HDAC7 lacking the first 22 amino acids, HDAC7-u^41^, in myeloid cells. In MacHDAC7 mice, *Csf1r* promoter elements direct myeloid-specific expression of the transcriptional activator GAL4-VP16, which in turn drives both HDAC7-u and cyan fluorescent protein (CFP) expression in myeloid cells. Littermate MacBlue mice that express GAL4-VP16-dependent CFP expression were used as controls in these studies^42^. We first evaluated the expression of inflammatory mediators and markers of liver fibrosis in naïve MacBlue and MacHDAC7 mice. As expected, mRNA levels of *Hdac7* were significantly elevated in the livers of MacHDAC7 mice compared to those of MacBlue control mice (**Figure 1A**). Thus, the effect of myeloid-specific *Hdac7* overexpression could be detected at the whole-organ level, even in the context of endogenous *Hdac7* expression in hepatocytes and other non-myeloid cells. Consistent with the elevated systemic levels of inflammatory mediators observed upon LPS challenge in MacHDAC7 mice^30^, mRNA levels of the inflammatory genes *Ccl2* and *Il1b* were increased in livers of these mice in a basal state (**Figure 1B-C**). Of the fibrotic genes that were examined, only *Mmp9* was increased (**Figure 1D**), with no difference being observed between the two genotypes in mRNA levels of other inflammatory or fibrotic genes such as *Crp*, *Fstl1*, *Col1a1* and *Act2* (**Figure S1**). Thus, myeloid *Hdac7* overexpression increases levels of specific inflammatory mediators in the liver, suggesting that it may prime liver inflammation and/or accelerate liver disease under certain conditions.

**Figure 1.**
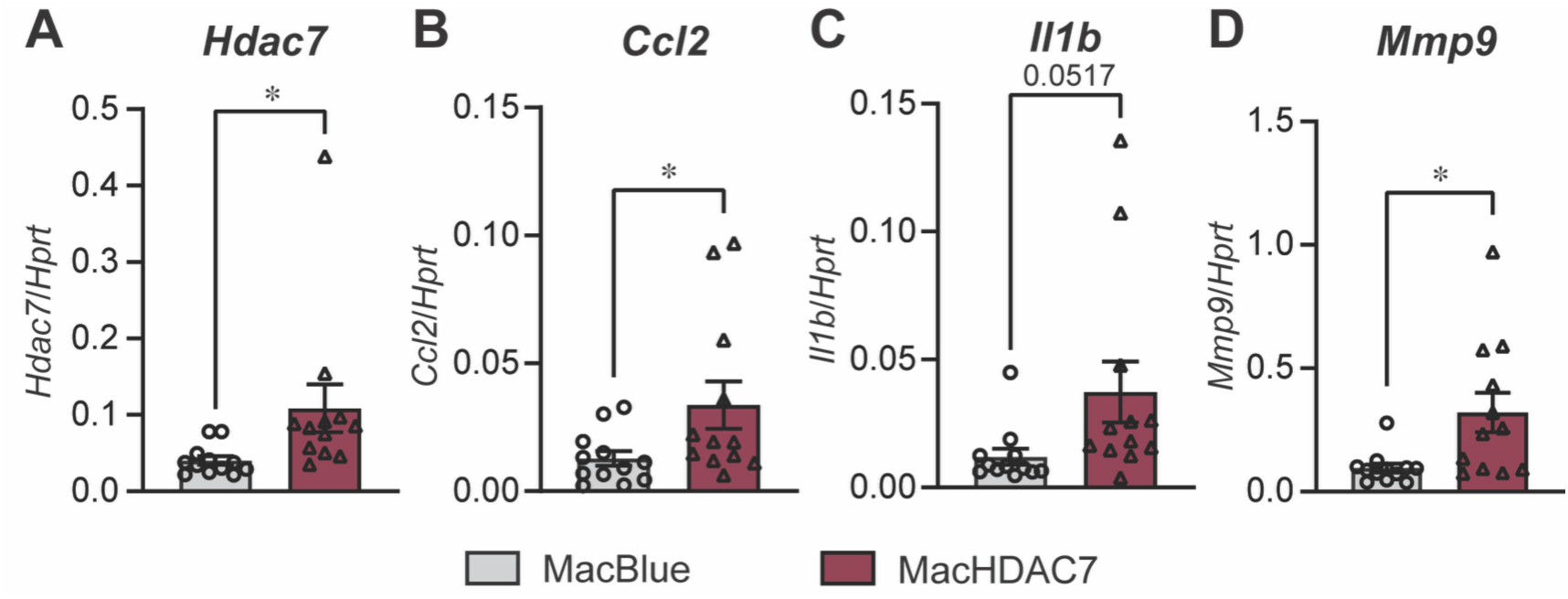
Myeloid *Hdac7* overexpression promotes expression of inflammatory genes in the liver. mRNA levels of *Hdac7* (A), *Ccl2* (B), *Il1b* (C) and *Mmp9* (D) in livers from 9-22 wk old MacHDAC7 and control MacBlue mice fed a chow diet. Data (normalised to *Hprt*) represent mean ± SEM from n=12 mice (6 female and 6 male) for each group. Statistical significance was determined by Student’s *t*-test (ns, not significant; * *p* <0.05).

### Myeloid *Hdac7* overexpression increases weight gain in mice fed a HFHCHS diet

To mimic the metabolic and histological changes of MASLD, we placed littermate MacHDAC7 and MacBlue control mice at 6-10 weeks of age on a HFHCHS diet (20% kcal fat, 20 g/kg cholesterol, 222 g/kg fructose and 107 g/kg sucrose) for 24 wk, with weekly body weight gain and metabolic parameters measured. As basal inflammatory mediators were elevated in livers of MacHDAC7 mice (**Figure 1**), cohorts of MacHDAC7 and MacBlue control mice fed on a chow diet were also included. Body weights of mice of each genotype were similar at the onset of the experiment (**Figure 2A**). As expected, the HFHCHS diet increased body weight for both genotypes by comparison to their respective chow diet groups. However, weight gain of MacHDAC7 mice fed on either a chow or HFHCHS diet was significantly reduced by comparison to the control MacBlue mice (**Figure 2A**). When correcting for this difference by normalizing the HFHCHS-dependent gain to the chow trajectory for each genotype, MacHDAC7 mice displayed a significantly greater relative weight gain on the HFHCHS diet than MacBlue over the 24 weeks of the experiment (**Figure 2B**). To study whole-body metabolism, fat and lean mass was also measured. Interestingly, *Hdac7* overexpression in myeloid cells reduced both the fat and lean mass in mice fed a chow diet (**Figure 2C**, left), consistent with the smaller average body weight of MacHDAC7 mice. This difference was not apparent in mice fed a HCHFHS diet (**Figure 2C**, right), suggesting that this diet increased both fat and lean body compositions in MacHDAC7 mice to a greater extent than MacBlue control mice.

**Figure 2.**
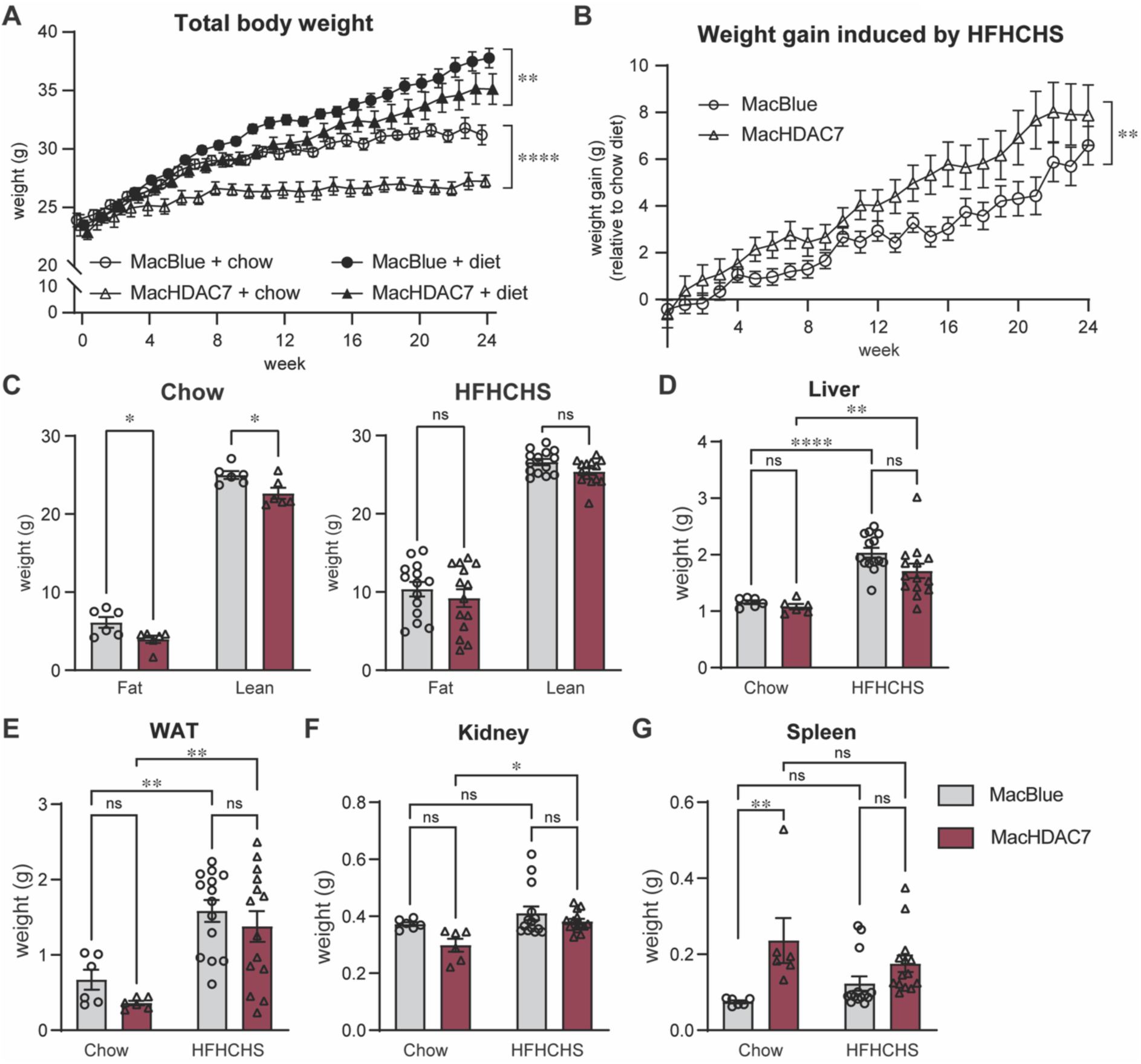
Effect of myeloid *Hdac7* overexpression on body weight gain in mice fed a chow versus HFHCHS diet. 8-10 wk old male MacBlue and MacHDAC7 mice were fed a chow diet (chow, n=6 per group) or a high fat, high cholesterol and high sucrose diet (HFHCHS, n=14-15 per group) for 24 wk. A. Weekly body weight. B. Average weight gain in mice fed a HFHCHS versus chow diet, calculated by substracting the average body weight of mice fed the chow diet from that of mice fed the HFHCHS. Data shown are mean ± SEM of n=6 (chow) or n=14-15 (HFHCHS) mice per group. Statistical analyses were perfomed using linear regression to compare the difference in body weight between MacBlue and MacHDAC7 mice fed on their respective diets or HFHCHS-induced weight gain (** *p* <0.01; **** *p* <0.0001). C. Fat and lean mass measured by EchoMRI at wk 23. D-G. Tissue weights of liver, epididymal white adipose tissue (WAT), kidney and spleen at wk 24. Data shown are mean ± SEM of n=6 (chow) or n=14 (HFHCHS) mice and were analysed using repeated measures (C) or ordinary (D-G) two-way ANOVA followed by Sidak’s multiple comparison (ns, not significant; * *p* <0.05; ** *p* <0.01; **** *p* <0.0001).

The HFHCHS diet also significantly increased the weights of liver and epididymal white adipose tissue (WAT) in both MacBlue and MacHDAC7 mice (**Figure 2D-E**), but this diet led to increased kidney weights only in MacHDAC7 mice (**Figure 2F**). This may reflect diet-induced renal damage. When normalising to the average tissue weight of chow-fed controls, the HFHSHC diet resulted in a modest, non-significant increase in epididymal fat weight for MacHDAC7 versus MacBlue mice (MacBlue 176.7% ± 46.3% vs. MacHDAC7 281.5% ± 56.3%, p = 0.07, Student’s t-test). In contrast, kidney weight was markedly elevated (MacBlue 9.87% ± 6.1% vs. MacHDAC7 27.0% ± 3.9%, p = 0.01, Student’s t-test). Interestingly, splenomegaly was observed in the MacHDAC7 mice when fed a chow diet (**Figure 2G**). This is suggestive of perturbed haematopoiesis in these mice, which would be consistent with *Hdac7*-induced liver inflammation in the basal state (**Figure 1**). We therefore next investigated the impact of HDAC7 overexpression on immune cell profiles. MacHDAC7 mice on the chow diet exhibited significant neutrophilia in peripheral blood, similar to levels observed for both MacBlue and MacHDAC7 mice on the HFHCHS diet (**Table 1**). This is consistent with a primed immune state that may drive low-grade myeloid inflammation associated with increased adiposity. Collectively, these data implicate myeloid *Hdac7* in both limiting weight gain of mice on a chow diet, as well enhancing relative weight gain of mice under HFHCHS. Both of these phenotypes could relate to HDAC7-driven inflammation and immune cell expansion.

**Table 1.** Haematological assessment of circulating immune cells in 30-34 week old MacBlue and MacHDAC7 mice fed a chow or HFHCHS diet.

|  | Chow |  | HFHCHS |  |
| --- | --- | --- | --- | --- |
|  | MacBlue | MacHDAC7 | MacBlue | MacHDAC7 |
| **WBC ($10^9/L$)** | $1.702 \pm 0.252$ | $1.783 \pm 0.211$ | $2.169 \pm 0.303$ | $1.835 \pm 0.26$ |
| **Neutrophil ($10^9/L$)** | $0.243 \pm 0.032$ | $0.725 \pm 0.157^a$ | $0.519 \pm 0.070$ | $0.634 \pm 0.123$ |
| **Monocyte ($10^9/L$)** | $0.045 \pm 0.007$ | $0.062 \pm 0.005$ | $0.079 \pm 0.009^b$ | $0.057 \pm 0.008$ |
| **Eosinophil ($10^9/L$)** | $0.027 \pm 0.008$ | $0.053 \pm 0.014$ | $0.031 \pm 0.005$ | $0.035 \pm 0.008$ |
| **Basophil ($10^9/L$)** | $0.003 \pm 0.002$ | $0.007 \pm 0.003$ | $0.005 \pm 0.002$ | $0.004 \pm 0.002$ |
| **Lymphoid ($10^9/L$)** | $1.383 \pm 0.218$ | $0.937 \pm 0.28$ | $1.535 \pm 0.249$ | $1.106 \pm 0.187$ |
| **RBC ($10^{12}/L$)** | $7.388 \pm 0.946$ | $7.93 \pm 0.279$ | $8.406 \pm 0.239$ | $8.187 \pm 0.153$ |
| **Platelet ($10^9/L$)** | $1128.833 \pm 186.582$ | $837.333 \pm 168.627$ | $906 \pm 80.965$ | $949.643 \pm 52.093$ |

Data shown are mean ± SEM and represent n =6 (chow) or n=14 (HFHCHS) mice for each genotype. Statistical analyses were performed using two-way ANOVA followed by Fisher's LSD test.
<sup>a</sup>MacBlue vs MacHDAC7 fed a chow diet, p=0.0230
<sup>b</sup>Chow vs HFHCHS in MacBlue group, p=0.0208

### Myeloid *Hdac7* overexpression drives neutrophilia and perturbs haematopoiesis

The immune profiling above was performed on mice aged 30-34 weeks, so it was conceivable that the observed neutrophilia was an age-assocated phenomenon. To investigate this, we assessed immune profiles of peripheral blood from 6-8 week old mice. Here we found that MacHDAC7 mice exhibited pronounced leukocytosis (p < 0.01) and neutrophilia (p < 0.05) compared with MacBlue controls, whereas there were no significant differences in monocytes, eosinophils, basophils and lymphocytes (**Table 2**). Young adult MacHDAC7 mice (6-10 week old) also showed marked splenomegaly versus MacBlue control mice (∼3-fold increase in spleen weight; p < 0.0001), with this accompanied by a substantial expansion of total splenocytes and splenic immune subsets, including B220⁺ B cells, CD3⁺ T cells, and CD11b⁺ myeloid cells (p < 0.0001) (**Table 3**). Thus, MacHDAC7 mice exhibit marked splenomegaly with broad immune-cell expansion and circulating leukocytosis dominated by neutrophils, indicative of a systemic immune-primed state.

**Table 2.**
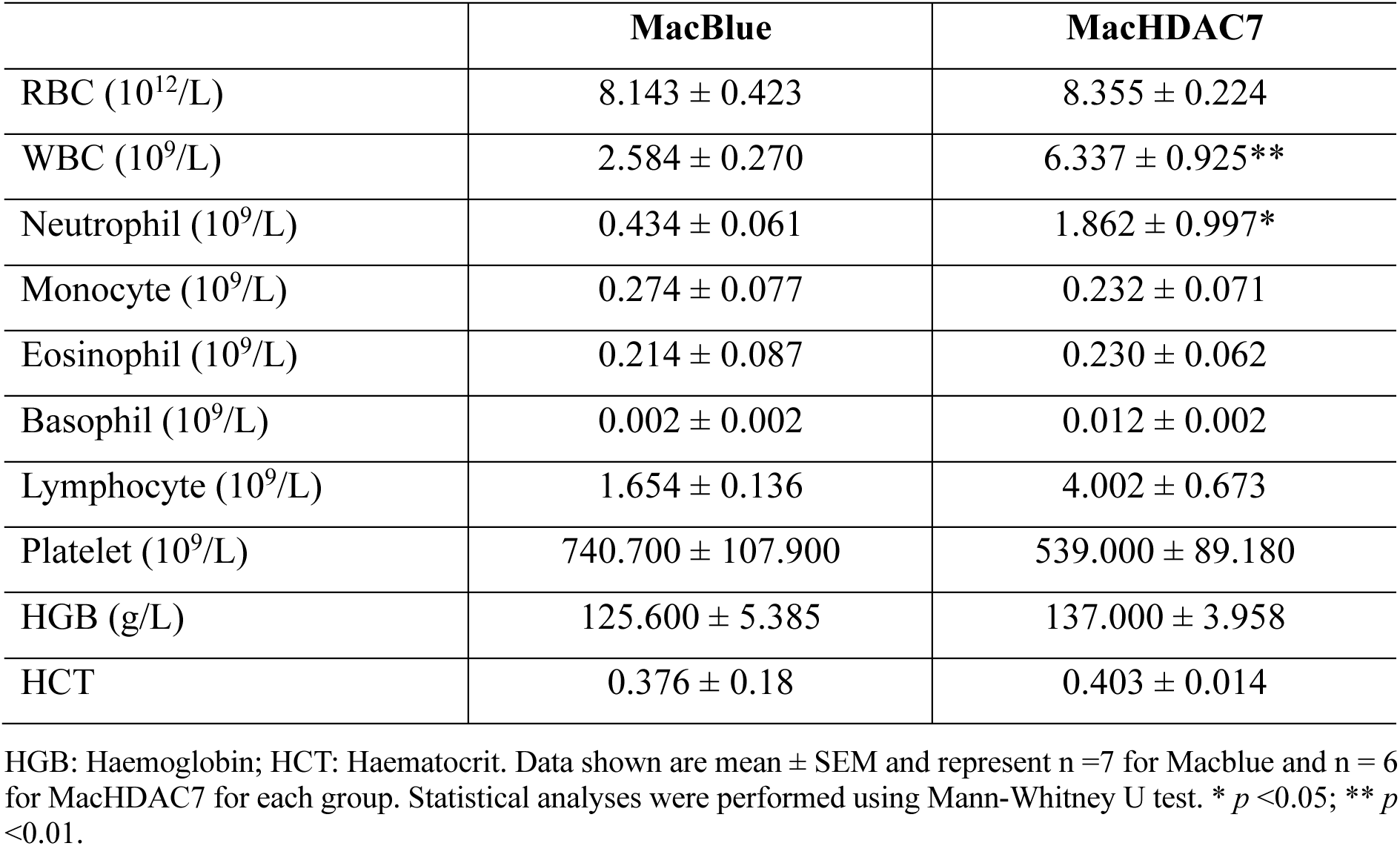
Haematological assessment of circulating immune cells in 6-8 week old MacBlue and MacHDAC7 mice.

|  | **MacBlue** | **MacHDAC7** |
| --- | --- | --- |
| RBC ($10^{12}/L$) | $8.143 \pm 0.423$ | $8.355 \pm 0.224$ |
| WBC ($10^9/L$) | $2.584 \pm 0.270$ | $6.337 \pm 0.925^{**}$ |
| Neutrophil ($10^9/L$) | $0.434 \pm 0.061$ | $1.862 \pm 0.997^*$ |
| Monocyte ($10^9/L$) | $0.274 \pm 0.077$ | $0.232 \pm 0.071$ |
| Eosinophil ($10^9/L$) | $0.214 \pm 0.087$ | $0.230 \pm 0.062$ |
| Basophil ($10^9/L$) | $0.002 \pm 0.002$ | $0.012 \pm 0.002$ |
| Lymphocyte ($10^9/L$) | $1.654 \pm 0.136$ | $4.002 \pm 0.673$ |
| Platelet ($10^9/L$) | $740.700 \pm 107.900$ | $539.000 \pm 89.180$ |
| HGB (g/L) | $125.600 \pm 5.385$ | $137.000 \pm 3.958$ |
| HCT | $0.376 \pm 0.18$ | $0.403 \pm 0.014$ |

HGB: Haemoglobin; HCT: Haematocrit. Data shown are mean ± SEM and represent n=7 for Macblue and n = 6 for MacHDAC7 for each group. Statistical analyses were performed using Mann-Whitney U test. \* $p < 0.05$ ; \*\* $p < 0.01$ .

**Table 3.** Assessment of spleen weight and immune cells in MacBlue and MacHDAC7.

|  | **MacBlue** | **MacHDAC7** |
| --- | --- | --- |
| Spleen weight (mg) | 73.000 ± 3.499 | 217.200 ± 19.760**** |
| Splenocytes (10<sup>6</sup> cells/spleen) | 86.210 ± 7.497 | 245.200 ± 27.79**** |
| B220+ B cells (10<sup>6</sup> cells/spleen) | 27.170 ± 2.957 | 70.270 ± 8.621**** |
| CD3+ T cells (10<sup>6</sup> cells/spleen) | 30.970 ± 4.840 | 47.800 ± 4.962* |
| CD11b+ Myeloid cells (10<sup>6</sup> cells/spleen) | 12.860 ± 2.720 | 83.720 ± 20.100**** |

Data shown are mean ± SEM and represent n = 10 for Macblue and n = 9 for MacHDAC7 for each group. Statistical analyses were performed using Mann-Whitney U test. \* $p < 0.05$ ; \*\*\*\* $p < 0.0001$ .

### Myeloid *Hdac7* promotes hepatic inflammation but not fibrosis in mice

Given that HDAC7 drives inflammatory responses in primary macrophages^30, 39^, we postulated that myeloid *Hdac7* overexpression alone may amplify diet-induced liver inflammation. As we previously observed (**Figure 1**), mRNA levels of *Ccl2* and *Il1b* were elevated in MacHDAC7 mice fed a chow diet compared to MacBlue control mice (**Figure 3A-B**). Interestingly, this difference was not apparent in mice fed a HFHCHS diet, suggesting that myeloid *Hdac7* alone primed liver inflammation. CCL2 and the accompanying infiltration of inflammatory monocytes contributes to MASH in animal models^43, 44^. We therefore assessed circulating CCL2 levels. MacHDAC7 mice fed a chow diet had significantly higher levels of serum CCL2 than the MacBlue control mice (**Figure 3C**). This effect was also apparent in mice fed the HFHCHS diet, although it was less pronounced because the HFHCHS diet increased CCL2 production in MacBlue mice. Consistent with its effects on hepatic *Ccl2* mRNA expression (**Figure 3A**) and circulating CCL2 (**Figure 3C**), hepatic mRNA levels of *Adgre1* that encodes the tissue macrophage marker F4/80 and Kupffer cell-restricted *Clec4f* were both elevated in MacHDAC7 mice versus MacBlue mice fed a chow diet (**Figure 3D-E**). Serum IL-1β levels were below the limits of detection in these samples (data not shown), but overall, the results suggest that myeloid *Hdac7* overexpression alone broadly phenocopies hepatic inflammation triggered by the HFHCHS diet. This was supported by blinded histological analyses, where liver inflammation was similar between MacHDAC7 mice on a chow diet and MacBlue control mice on a HFHCHS diet (**Figure 3F-G**).

**Figure 3.**
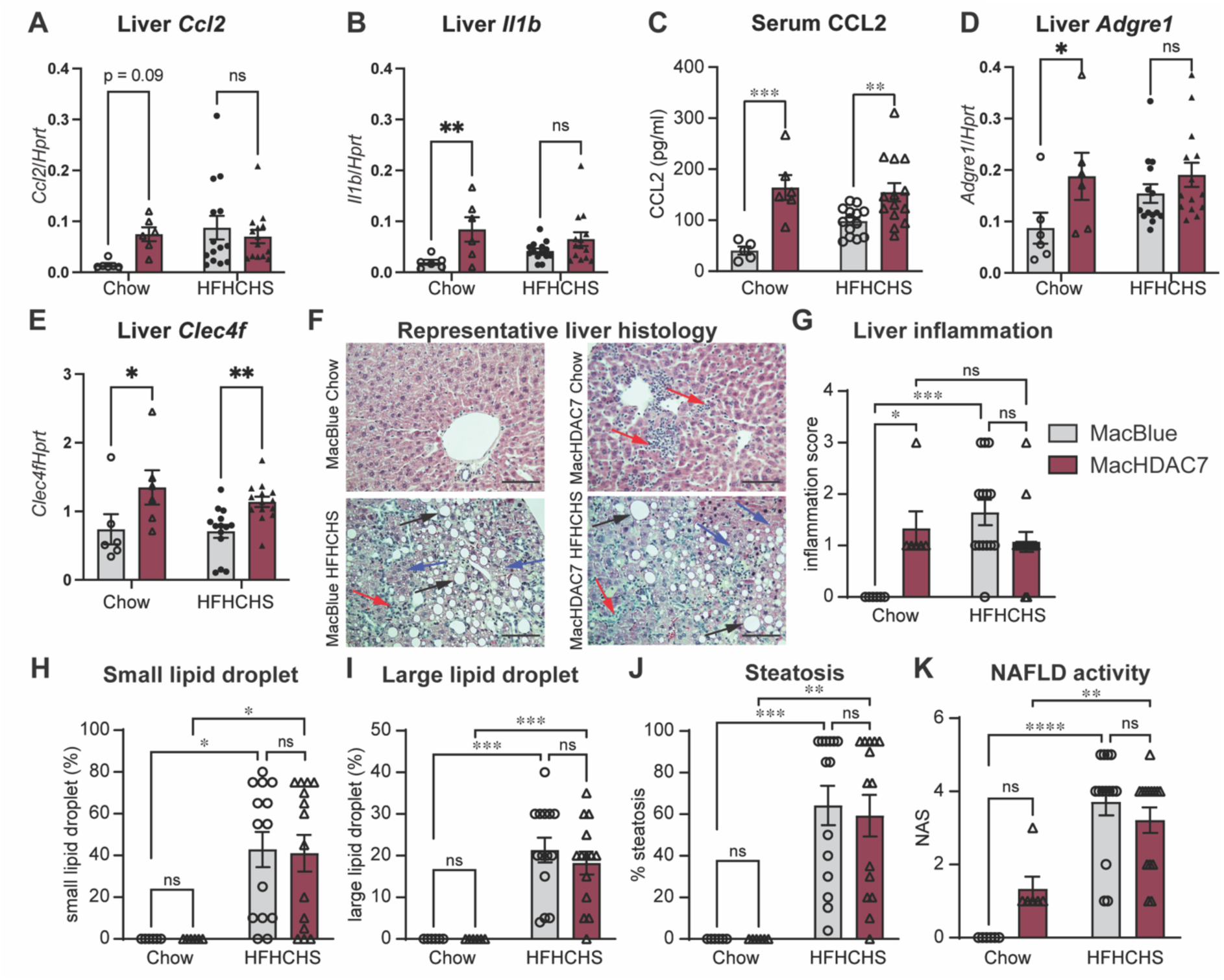
Myeloid *Hdac7* overexpression drives hepatic inflammation but not steatosis. Hepatic mRNA levels of genes encoding inflammatory mediators and serum CCL2 levels in male MacBlue and MacHDAC7 mice fed a chow or HFHCHS diet. A-B. Liver mRNA levels of *Ccl2* (A) and *Il1b* (B) relative to *Hprt* were measured by RT-qPCR. C. CCL2 levels in mouse serum were measured by ELISA. D-E. Liver mRNA levels of F4/80-encoding *Adgre1* (D) and *Clec4f* (E) relative to *Hprt* were measured by RT-qPCR. F. Representative H&E staining of livers from chow- or HFHCHS-fed MacBlue and MacHDAC7 mice. Photos were acquired at 40 x magnification, scale bar: 100 μm. Large lipid droplets (black arrows) and small lipid droplets (blue arrows) and a focus of inflammation within the lobule (red arrows). G-I. Quantification of lobular inflammation (G), lipid droplet counts (H-I), steatosis (J) and NAFLD activity score (K), as per the NASH clinical research network scoring system after H&E staining. Data represent mean ± SEM of n=6 (chow) or n=14 (HFHCHS) mice for each treatment group and were analysed using two-way ANOVA followed by Fisher’s LSD test (A-E) or Sidak’s multiple comparison (G-K) (ns, not significant; * *p*< 0.05; ** *p*< 0.01; *** *p*< 0.001; **** *p*< 0.0001).

Having established a role for myeloid HDAC7 in driving liver inflammation, we next assessed its role in liver steatosis and fibrosis. Here we found that the HFHCHS diet induced lipid accumulation and liver steatosis in both MacBlue and MacHDAC7 mice, but this effect was not amplified by *Hdac7* overexpression (**Figure 3H-J**). This suggests that myeloid *Hdac7* overexpression does not have obvious effects on lipid metabolism in the liver. Although histological examination revealed that myeloid *Hdac7* overexpression did not exacerbate HFHCHS-induced liver inflammation (**Figure 3F-G**), we did observe a modest and non-significant increase in NAFLD activity score in MacHDAC7 mice fed a chow diet by comparison to versus MacBlue control mice (**Figure 3K**). However, the HFHCHS diet failed to induce a constant and robust MASH-like phenotype in either group, as shown by the mild and highly variable levels of fibrosis (**Table S1**), thus it remains unclear whether myeloid *Hdac7* promotes diet-induced MASH. Taken together, these observations suggest that myeloid *Hdac7* overexpression promotes hepatic and systemic inflammation, but this alone is not sufficient to drive liver fibrosis.

### Glucose metabolism is dysregulated in MacHDAC7 mice fed a HFHCHS diet

Insulin resistance is a common metabolic syndrome of MASLD^45^. Thus, we measured fasted blood glucose levels to assess whether myeloid *Hdac7* overexpression regulates hepatic glucose metabolism. When placed on a chow diet, MacHDAC7 mice consistently had lower fasted blood glucose levels than the control MacBlue mice, with these effects being significant at wk 4 and 16 (**Figure 4A**). To assess glucose clearance, we performed intraperitoneal glucose tolerance tests (GTTs) at wk 12, finding that the MacHDAC7 mice cleared injected glucose more rapidly than the MacBlue control, albeit the difference was not statistically significant (**Figure 4B**). However, these phenotypes were reversed when mice were fed a HFHCHS diet, with higher blood glucose levels and slower glucose clearance observed in MacHDAC7 mice (**Figure 4C-D**). This suggests that myeloid HDAC7 has context-dependent roles in regulating glucose metabolism.

**Figure 4.**
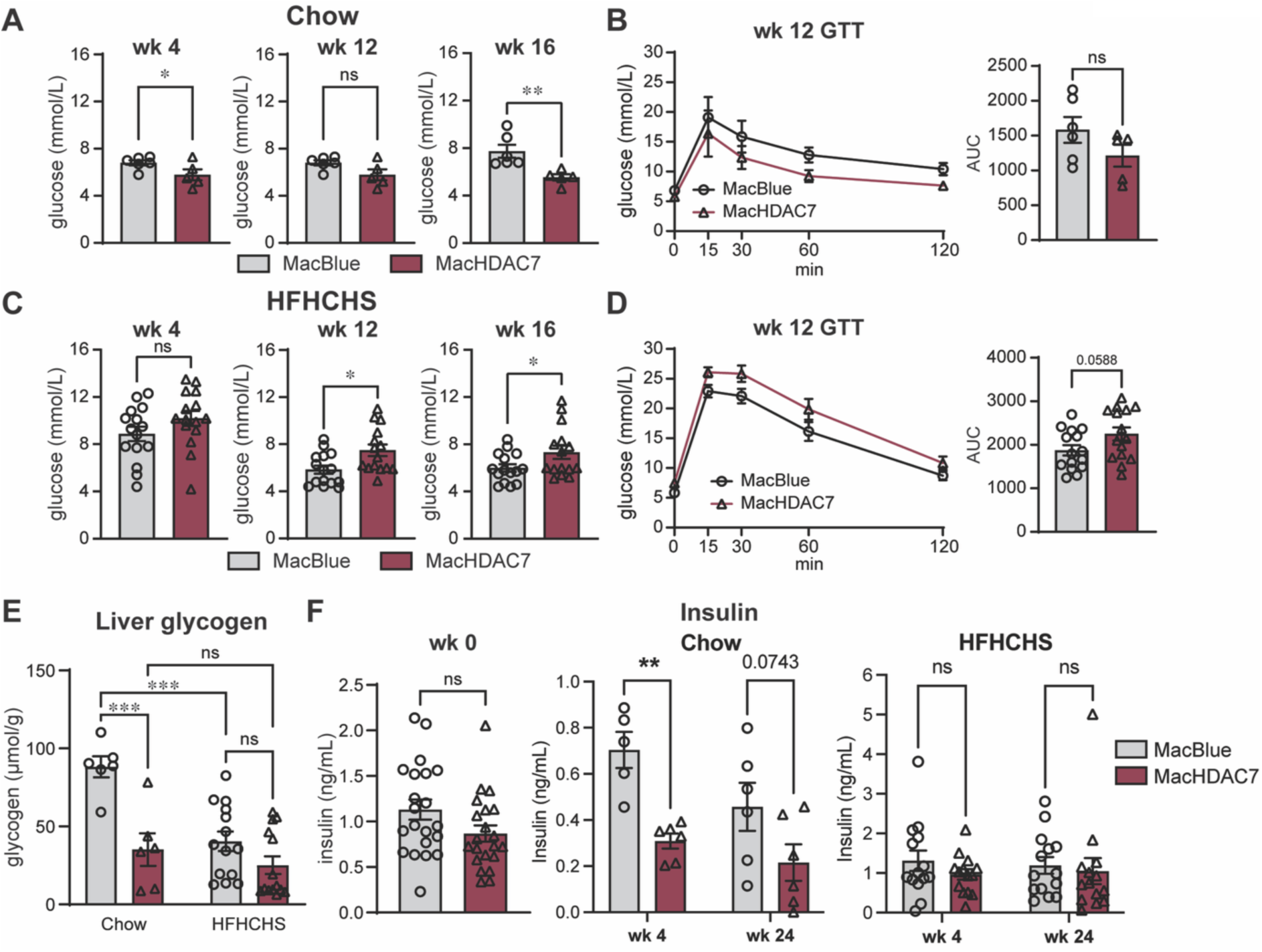
Myeloid *Hdac7* overexpression increases fasted blood glucose levels, glucose dysregulation and glycogen metabolism in mice fed a HFHCHS diet. 6 h fasted blood glucose levels of MacBlue and MacHDAC7 mice fed a chow (**A**) or HFHCHS (**C**) diet. Glucose tolerance test (GTT) was performed at wk 12 (**B** and **D**). Data shown are mean ± SEM of n=5-6 (chow) or n=15 (HFHCHS) mice for each group. AUC, area under curve analysis. **E.** Liver glycogen quantification. At termination of the study, mice were fasted for 6 h and glycogen levels were quantified in snap-frozen livers. Data (mean ± SEM) represent n=6 (chow) or n=14 (HFHCHS) mice for each group. **F.** Insulin levels in the circulation. Circulating insulin levels were measured before diet feeding (wk 0), at wk 4 and at end-point, after mice were fasted for 6 h. Data are mean ± SEM of n=5-6 (chow) or n=14 (HFHCHS) mice for each treatment group. Statistical analyses were performed using Student’s *t*-test (**A-D, F** wk 0) or two-way ANOVA followed by Sidak’s multiple comparison (**B, D, E-F**) (ns, not significant; * *p*< 0.05; ** *p*< 0.01; *** *p*< 0.001).

In a fasting state, glucose is mainly produced in the liver via glycogenolysis and gluconeogenesis, with the former converting glycogen into glucose^46^. We therefore considered that hyperglycaemia in the MacHDAC7 mice fed a HFHCHS diet could be due to a change in hepatic glycogen metabolism. Interestingly, MacHDAC7 mice had lower liver glycogen levels than MacBlue control mice when fed the chow diet, with a similar non-significant trend being apparent for mice fed the HFHCHS diet (**Figure 4E**). It is therefore likely that myeloid HDAC7 plays a role in glycogen metabolism in both lean and obese mice. Hyperglycaemia inhibits glycogenolysis in healthy individuals, with this inhibition being impaired in type 2 diabetes mellitus^47^. Since MacHDAC7 mice fed a HFHCHS diet had lower hepatic glycogen levels (**Figure 4E**) and increased fasting glucose levels (**Figure 4C**) than MacBlue mice, we propose that regulatory control of hepatic glycogen breakdown may be impaired in MacHDAC7 mice, leading to enhanced glycogenolysis. We conclude that myeloid HDAC7 overexpression exacerbates HFHCHS diet-induced dysregulation of glucose and glycogen metabolism.

Inflammatory mediators such as CCL2 promote macrophage infiltration, hepatic inflammation, and insulin resistance during diet-induced obesity^48, 49^. Considering the role of HDAC7 in macrophage inflammation and that myeloid *Hdac7* expression triggered higher levels of circulating CCL2 (**Figure 3C**), we next wondered whether myeloid *Hdac7* alters insulin secretion during obesity. Indeed, insulin levels were reduced in MacHDAC7 mice versus MacBlue control mice fed a chow diet (**Figure 4F**, centre). This suggests that lean MacHDAC7 mice (**Figure 2C**) control insulin levels to respond to their lower blood glucose levels (**Figure 4A**). Interestingly, when fed a HFHCHS diet, both MacBlue and MacHDAC7 mice had similarly elevated insulin levels (**Figure 4F**, right), suggesting a feedback mechanism of increased insulin production to suppress elevated blood glucose levels. However, these increased insulin levels failed to control glucose production in MacHDAC7 mice (**Figure 4C**). We conclude that elevated HDAC7 expression in myeloid cells drives metabolic dysregulation, a feature of type 2 diabetes mellitus^50^.

### Myeloid *Hdac7* deletion does not alter weight gain but attenuates HFHCHS diet-induced glucose dysregulation

Given that myeloid *Hdac7* overexpression triggered neutrophilia and hepatic inflammation, as well as dysregulated glucose homeostasis in diet-induced obesity, we hypothesised that genetically targeting *Hdac7* in myeloid cells would ameliorate HFHCHS-induced phenotypes. To test this, we used myeloid-restricted *Hdac7*^-/-^ mice (*Hdac7*^fl/fl^ *LysM*^Cre^) and littermate *Hdac7*^+/+^ control mice (*Hdac7*^fl/fl^), placing the mice on a HFHCHS diet for a period of 36 wk with the goal of inducing more severe MASH phenotypes. In contrast to the increased relative weight gain of myeloid HDAC7 gain-of-function mice on a HFHCHS versus chow diet (**Figure 2B**), *Hdac7* deletion from myeloid cells did not affect weight gain in mice fed a HFHCHS diet (**Figure S2A**). Similarly, fat and lean mass (**Figure S2B**) and organ weights (**Figure S2C**) were unaffected, although there was a modest trend towards reduced liver mass. These data indicate that myeloid *Hdac7* deficiency does not prevent the development of obesity.

Next, we examined glucose metabolism to determine whether myeloid *Hdac7* deletion improves the dysregulated phenotypes induced by the HFHCHS diet. *Hdac7*^-/-^ mice had similar blood glucose levels to the control mice at the beginning of the study (**Figure 5A**, 0 wk). However, starting from wk 8 of HFHCHS feeding, myeloid-restricted *Hdac7*^-/-^ mice had consistently lower fasted blood glucose levels than *Hdac7*^+/+^ mice, with the effects being significant at wk 8, 14 and 16 (**Figure 5A**). This suggests that deletion of *Hdac7* from myeloid cells improves blood glucose regulation. This effect was also reflected in improved glucose tolerance in *Hdac7*^-/-^ mice (**Figure 5B-C**). *Hdac7*^-/-^ mice also presented with slightly improved insulin sensitivity than control mice, although these effects were not significant (**Figure 5D-E**). Hence, myeloid deletion of *Hdac7* reduces fasting glucose and improves glucose intolerance in the HFHCHS obesity model.

**Figure 5.**
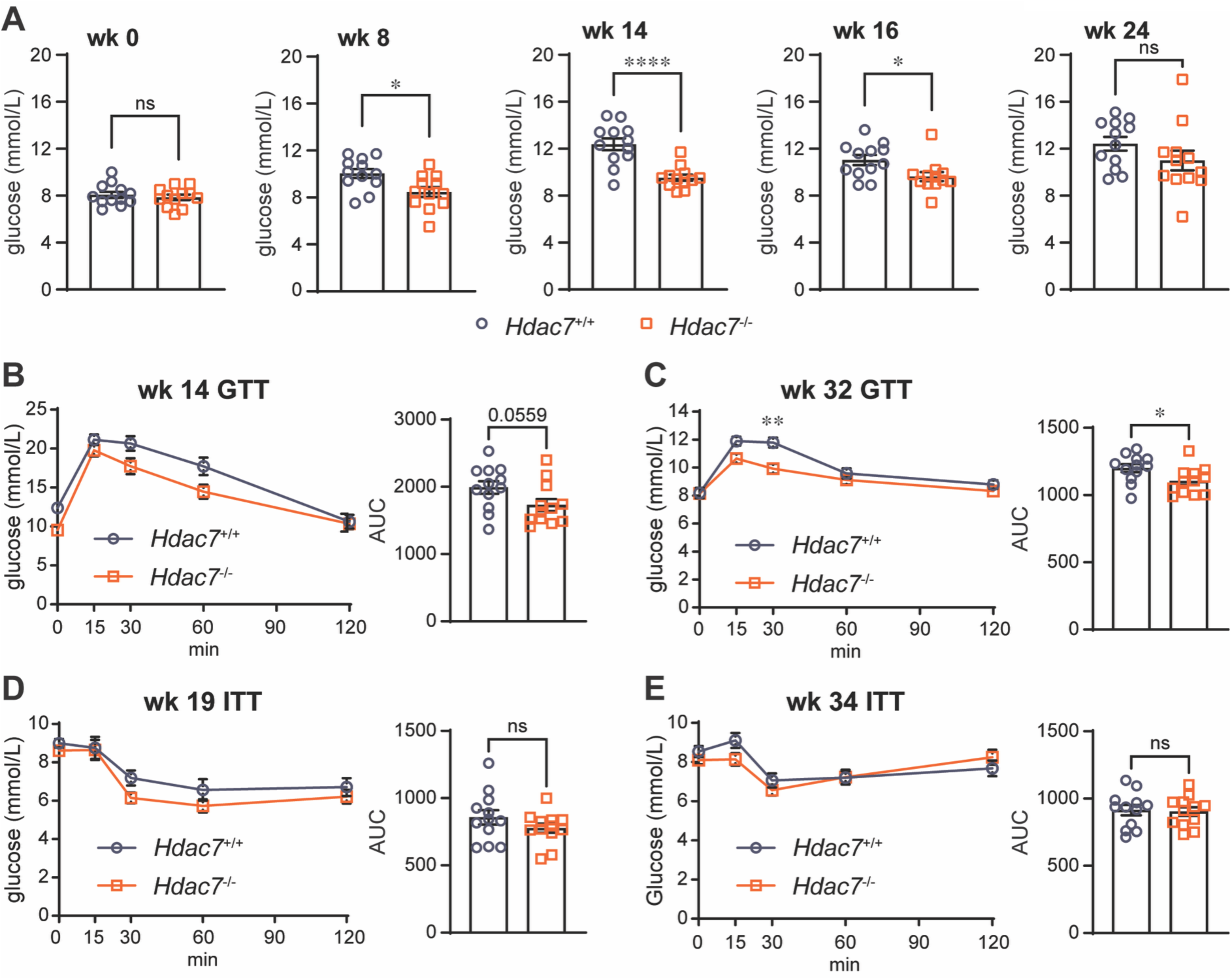
Myeloid *Hdac7* deficiency reduced fasting blood glucose levels in mice fed a HFHCHS diet. 8-10 wk old male *Hdac7*^+/+^ and *Hdac7*^-/-^ mice were fed a HFHCHS diet for 36 wk. **A.** Fasted blood glucose levels were measured at wk 0, 8, 14, 16 and 24. Data are mean ± SEM of n=12 mice for each group and were analysed using Student’s *t*-test (ns, not significant; * *p* < 0.05; **** *p* < 0.0001). Glucose tolerance test (GTT) at wk 14 and wk 32 (**B-C**) and insulin tolerance test (ITT) at wk 19 and wk 34 (**D-E**). Data shown are mean ± SEM of n=12 mice/ group. Statistical analyses were performed using repeated measures two-way ANOVA followed by Sidak’s multiple comparison. Area under curve (AUC) of GTTs and ITTs were analysed using Student’s *t*-test (ns, not significant; * *p* < 0.05; ** *p* < 0.01).

### Myeloid *Hdac7* deletion did not ameliorate diet-induced hepatic pathology and inflammation

Next, we assessed whether the effects of *Hdac7* deletion on glucose metabolism may relate to inflammation. At the cellular level, levels of HDAC7 protein were greatly reduced in bone marrow derived macrophages (BMMs) from *Hdac7*^-/-^ mice fed on both HFHCHS and chow diets, as expected (**Figure S3A**). Consistent with previous findings^30^, LPS-inducible CCL2 and inflammasome-triggered IL-1β levels were reduced in BMMs from the naïve *Hdac7*^-/-^ mice on a chow diet (**Figure S3B**, right side of graphs). However, in *Hdac7*^-/-^ BMMs generated from mice fed the HFHCHS diet, only IL-1β levels were significantly reduced, with CCL2 levels being only marginally affected (**Figure S3B**, left side of graphs). These data indicate that the capacity of HDAC7 to drive the production of these inflammatory mediators in macrophages may be attenuated under the conditions of obesity. Consistent with this, hepatic *Ccl2* mRNA levels were only slightly reduced in myeloid *Hdac7*^-/-^ mice (**Figure S3C**). Liver *Il1b* mRNA levels (**Figure S3D**) and circulating CCL2 levels (**Figure S3E**) were also unaffected. Histological assessment of livers from these mice revealed that the HFHCHS diet induced only varying levels of early to moderate stages of liver fibrosis in mice from either genotype (**Figure S4A**). Of note, the *Hdac7*^-/-^ mice trended towards a very modest reduction in liver inflammation, lipid accumulation, steatosis and fibrosis compared to the *Hdac7*^+/+^ control mice (**Figure S4B-F**). This suggests that myeloid *Hdac7* deletion may very modestly reduce obesity-induced pathological processes in the liver. We conclude that myeloid HDAC7 drives glucose dysregulation but only marginally contributes to hepatic inflammation and fibrosis in the HFHCHS diet model.

### Myeloid *HDAC7* expression is associated with gene expression signatures linked to advanced CLD

Our data support a role for HDAC7 in mediating glucose dysregulation in both gain- and -loss of function mouse models, as well as liver inflammation in a gain-of-function mouse model. We therefore considered that elevated HDAC7 expression in CLD may contribute to metabolic dysregulation and/or disease progression. We thus assessed *HDAC7* mRNA levels in liver biopsies of people with early or advanced CLD^51^. Here we found that *HDAC7* mRNA levels were elevated in livers from people with advanced CLD by comparison to those with early disease (**Figure 6A**), suggestive of a link between HDAC7 and human CLD severity. To further investigate this link in myeloid cells, we performed spatial transcriptomic analysis on livers from MacBlue and MacHDAC7 mice fed a chow diet to identify myeloid HDAC7-associated transcriptional changes relevant to CLD. We first identified cells co-expressing *eCFP* and *Hdac7* mRNAs in livers, then performed differential gene expression analysis on *eCFP*^+^/*Hdac7*^+^ cells from MacBlue versus MacHDAC7 mice. A heatmap of the 53 differentially expressed genes is shown in **Figure 6B**. This analysis confirmed elevated *Il1b*, *Ccl2* and *Mmp9* mRNA levels in hepatic myeloid cells from MacHDAC7 mice (**Figure 6B**), consistent with our previous findings in whole liver tissue (**Figure 1B-D**). When compared with our previously published dataset of genes associated with early and advanced human CLD^51^, we found that eight genes associated with advanced CLD in human were also differentially expressed in *eCFP*^+^ cells in livers of MacHDAC7 mice. These include *Vcan*, *Thbs1*, *Mmp9* and *Rab8b* (**Figure 6B**), several of which have been implicated in CLD progression and/or pathology^52-54^. Spatial co-expression analysis revealed that these genes were co-expressed with HDAC7, with limited expression in other hepatic cells (**Figure 6C-F**). This suggests that myeloid HDAC7 expression is sufficient to direct gene expression programs associated with advanced CLD. In summary, data presented in this study implicate myeloid HDAC7 in driving hepatic inflammation and expression of genes linked to advanced CLD, as well as systemic glucose dysregulation in a mouse obesity model.

**Figure 6.**
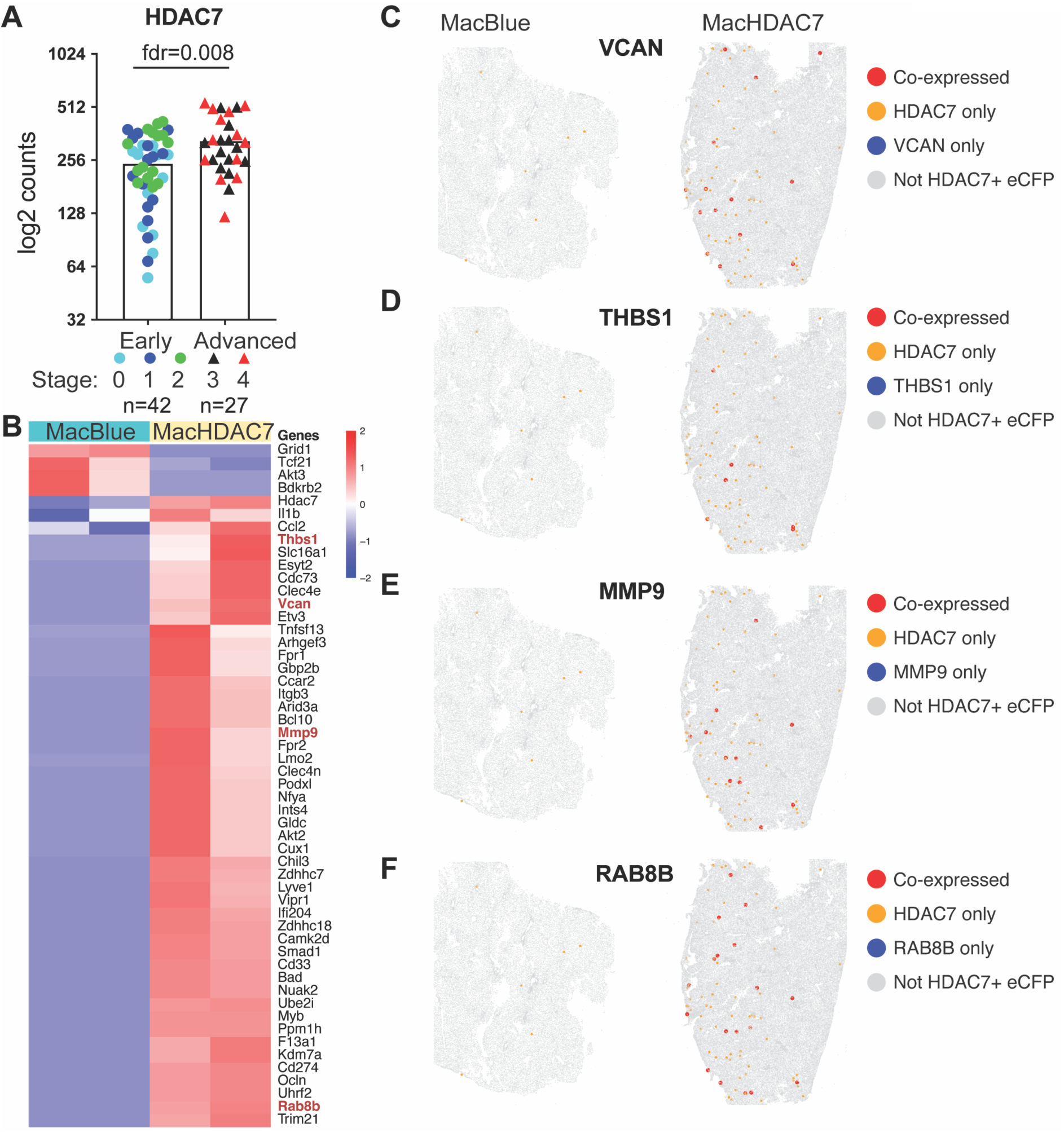
Association of *HDAC7* mRNA expression with signatures of human CLD. **A.** mRNA levels of *HDAC7* in liver biopsies from patients with early (stages 0-2) versus advanced (stages 3-4) of CLD was determined by RNAseq. Differentially expressed genes were identified using edgeR with TMM normalisation and Benjamini-Hochberg method for fdr correction. Data indicate mean ± SEM, n = 69 (circles = early stages and triangles = advanced stages). **B-F.** Spatial transcriptomic profiling of liver tissues from MacBlue and MacHDAC7 mice was performed using the 10x Genomics Xenium platform. Expression of eCFP and HDAC7 mRNAs was used to identify hepatic myeloid cells expressing HDAC7 (CFP⁺/HDAC7⁺ cells). **B.** A heatmap showing the 53 differentially expressed genes in CFP⁺/HDAC7⁺ macrophages from MacBlue versus MacHDAC7 livers. **C-F.** Spatial co-expression analysis of *VCAN* (**C**), *THBS1* (**D**), *MMP9* (**E**) and *RAB8B* (**F**) in MacBlue and MacHDAC7 livers. Data are representative of two independent tissue samples from litter-matched male MacBlue and MacHDAC7 mice fed a chow diet.

## Discussion

In this study, we investigated potential roles of myeloid HDAC7 in metabolic diseases, examining obesity, glucose dysregulation, hepatic inflammation and fibrosis. Although levels of steatosis and fibrosis were variable in our studies (**Figure 3H-K, S4C-F, Table S1**), HFHCHS fed mice did develop diet-induced obesity (**Figure 2A**) and liver inflammation (**Figure 3F-G**). We therefore focused our studies on indicators of metabolism and inflammation. We found the overexpression of myeloid *Hdac7* promoted hepatic inflammation (**Figure 3**) and expansion of leukocytes, particularly neutrophils (**Tables 1-2**), in a basal state, sharing some overlap with phenotypes observed in control MacBlue mice under HFHCHS. We also observed that weight gain of MacHDAC7 mice fed either a chow or HFHCHS diet was reduced by comparison to MacBlue mice (**Figure 2A**), which may relate to the energetic burden required for immune cell expansion. These findings complicate interpretation of the effects of HDAC7 on HFHCHS diet-induced weight gain, though we note that the relative increase in body weight (HFHCHS versus chow diet) was significantly increased in MacHDAC7 versus MacBlue mice (**Figure 2B**). Of note, in the HFHCHS diet model, glucose dysregulation was exacerbated in myeloid HDAC7 gain-of-function mice and improved in myeloid HDAC7 loss-of-function mice. This key finding thus extends the functions of HDAC7 in macrophages beyond cellular metabolism^30, 39, 40^ to whole body glucose metabolism.

MacHDAC7 mice on a HFHCHS diet had increased hyperglycaemia at multiple time points (**Figure 4C**), as well as impaired glucose tolerance (**Figure 4D**), by comparison to MacBlue control mice. Conversely, hyperglycaemia (**Figure 5A**) and glucose clearance (**Figure 5B**) were improved in myeloid *Hdac7*^-/-^ mice on a HFHCHS diet. Collectively, these data implicate myeloid HDAC7 as a contributor to glucose dysregulation during obesity. Interestingly, MacHDAC7 mice on a chow diet had reduced levels of fasted blood glucose (**Figure 4A**) and liver glycogen (**Figure 4E**), suggesting that glycogen metabolism is disrupted in these mice. Under fasting conditions, glycogenolysis is initiated by glucagon and epinephrine-dependent cascades^55^. Glycogen phosphorylase is a key enzyme controlling glycogenolysis, converting glycogen to glucose-1-phosphate, which is subsequently converted to glucose-6-phosphate that feeds glycolysis^56^. A previous study found that glycogen phosphorylase activity is controlled by post-translational modifications. Specifically, acetylation at K470 inhibits its enzymatic activity, with this inactivation mechanism being enhanced by insulin^57^. MacHDAC7 mice on the chow diet had significantly lower blood insulin levels than control MacBlue mice, with this difference being absent in mice fed the HFHCHS diet (**Figure 4F**). Thus, we propose that myeloid HDAC7 may somehow limit insulin secretion in MacHDAC7 mice to indirectly enhance glycogenolysis in hepatic cells. This hypothesis will need to be tested in future studies to characterise hepatic glycogen content and flux, as well as molecular mechanisms involved. Alternatively, it is also possible that HDAC7 promotes glycogen breakdown in macrophages, which could contribute to glucose dysregulation in the setting of obesity. Activated macrophages have been shown to store glycogen and metabolise it to drive inflammatory responses via multiple mechanisms^58^. Thus, HDAC7-mediated glycogenolysis in macrophages could contribute to both dysregulated inflammation and glucose metabolism during obesity. We previously showed that HDAC7 increases the enzymatic activity of the glycolytic enzyme PKM2 by interacting with it and promoting its deacetylation^30^ and it is possible that HDAC7 similarly activates enzymes in the glycogenolysis pathway in macrophages.

In the fasting state, another source of hepatic glucose production is gluconeogenesis, which is under strict hormonal control. Silencing of HDAC4, 5 and 7 in mouse livers led to higher liver glycogen content and improved glucose tolerance^59^. Mechanistically, the fasting hormone glucagon promotes the dephosphorylation and nuclear shuttling of class IIa HDACs in HepG2 hepatocytes. In the nucleus, class IIa HDACs recruit HDAC3 to promoter regions of gluconeogenic genes to enable the deacetylation of the FOXO transcription factor and the induction of downstream genes, including *G6pc* that encodes glucose-6-phosphatase, a key enzyme mediating gluconeogenesis^59^. An analogous pathway of hormone-activated HDAC4/FOXO regulation of glucose and lipid metabolism was also identified in *Drosophila melanogaster*^60^. Given these studies, it is possible that myeloid HDAC7 also contributes to hyperglycaemia via activating gluconeogenic genes downstream of glucagon signalling.

Liver inflammation in MacHDAC7 and MacBlue mice on the HFHCHS was comparable (**Figure 3F-G**), in contrast to the increased inflammation in MacHDAC7 mice on a chow diet (**Figure 3A-G**). The lack of an enhanced inflammatory phenotype in obese MacHDAC7 mice may reflect that HDAC7 overexpression could share some features of diet-induced inflammation. In contrast, levels of inflammatory mediators were not significantly reduced in the livers or plasma of myeloid *Hdac7*^-/-^ mice fed a HFHCHS diet (**Figure S3-S4**), although BMMs isolated from these mice on a chow diet showed reduced production of inflammatory mediators (**Figure S3B**, right side of graphs). The lack of an obesity-driven inflammatory phenotype in myeloid *Hdac7*^-/-^ mice may reflect compensatory effect of other class IIa HDACs, as previously observed in activated macrophages *in vitro*^30^. It may also reflect variability in efficiency and/or specificity of the LysM-Cre system for conditional gene deletion in myeloid cells^61^, as highlighted by a previous study examining immune and non-immune cell populations of the lung^62^. Alternatively, myeloid HDAC7 may have distinct roles in obesity- versus LPS-driven inflammation. We previously showed that HDAC7 promotes glycolysis and associated IL-1β production in macrophages responding to LPS^30^. However, during bacterial infection, macrophage HDAC7 engages the pentose phosphate pathway to generate the metabolite ribulose-5-phosphate that suppresses macrophage inflammatory responses^40^. This context-dependent role of myeloid HDAC7 in controlling macrophage inflammatory responses makes it difficult to predict its functions in different inflammation-driven conditions. Nonetheless, taken together with the pronounced effects of myeloid gain-of-function HDAC7 in exacerbating hepatic inflammation (**Figure 1, 3**), the subtle reduction in phenotypes associated with MASLD in myeloid *Hdac7*^-/-^ mice (**Figure S4**) is suggestive of a potential role for HDAC7 in contributing to hepatic inflammation during metabolic dysfunction. The involvement of myeloid HDAC7 in glucose dysregulation, as demonstrated through both gain-of-function (**Figure 4**) and loss-of-function (**Figure 5**) models, is consistent with such a link.

In conclusion, this study has identified an association between myeloid HDAC7 and both hepatic and systemic glucose metabolism, thus identifying a potential role for macrophage-expressed HDAC7 in controlling whole body metabolism. This lysine deacetylase and scaffolding protein likely promotes the breakdown of liver glycogen, which may partly explain HDAC7-mediated hyperglycaemia during obesity. Future studies should address molecular mechanisms by which HDAC7 mediates glycogenolysis and systemic glucose dysregulation. Myeloid HDAC7 overexpression also increased hepatic inflammation in mice under a chow diet, but an important limitation of our study is that we did not identify the mechanisms involved. Nonetheless, our findings are consistent with the known role of HDAC7 in driving both PKM2-mediated inflammatory responses^30, 39, 63^ and oxidative stress^40^ in macrophages. In contrast, although myeloid *Hdac7* deletion improved glucose regulation under HFHCHS, it did not reduce hepatic inflammation. Thus, endogenous HDAC7 is not essential for driving diet-induced liver inflammation. Nonetheless, *HDAC7* expression was elevated in liver biopsies of people with advanced versus early CLD (**Figure 6A**) and in tumours derived from a mouse liver carcinoma model^64^, with myeloid HDAC7 directing expression of genes associated with advanced CLD in human (**Figure 6B-F**). Thus, it will be of interest in the future to evaluate whether small molecule inhibitors of class IIa HDACs^65, 66^ can attenuate dysregulated metabolism, chronic inflammation and/or liver fibrosis in mouse models of obesity-driven CLD.

## Materials and methods

### Ethics statement

All animal studies were approved by The University of Queensland Animal Ethics Committee. Experimental procedures followed the approved projects 2021/AE00629, 2021/AE00630 and 2021/AE001048. Mice were bred and housed on a 12 h-controlled day/night cycle with food and water available ad libitum in the Queensland Biosciences Precinct animal house (Institute for Molecular Bioscience, The University of Queensland). Mice were monitored during experiments in accordance with protocols approved in the above animal ethics certificates.

### Animals

6-10 wk old male MacHDAC7 mice and control MacBlue mice (total of n=14-15 per group) were fed a HFHCHS diet containing 20% kcal fat, 20 g/kg cholesterol, 222 g/kg fructose and 107 g/kg sucrose (Specialty Feeds, SF21-160, containing sodium caseinate) for 24 wk. The control groups (n=3 per genotype) were fed a chow diet for the same time. These experiments were carried out as two independent trials. Alternatively, 8-12 wk old male *Hdac7*^-/-^ and wild type control *Hdac7*^+/+^ mice (n=12 per genotype) were fed a HFHCHS diet (Specialty Feeds, SF18-074, same nutrient composition to SF21-160 except that it contains casein) for 36 wk. Body weight was measured weekly throughout the trials. Fat and lean mass was measured using a Minispec LF50 NMR Analyser (Bruker) one week before the end of the study. At the termination of the study, mice were fasted for 6 or 12 h before being euthanised with isoflurane. Blood was collected via cardiac puncture followed by perfusion with 0.9% saline solution (Baxter). Liver, spleen, kidney and epididymal white adipose tissue were collected, weighed, snap frozen, then fixed in 10% neutral buffered formalin or processed for RNA extraction. Relative tissue weight gain for mice of each genotype was calculated by normalizing each HFHCHS-fed mouse to the mean chow-fed tissue weight using the formula: [(HFHCHS tissue weight − chow mean)/chow mean] × 100%.

### Fasted glucose measurements

Mice were fasted for 6 h before blood was collected at midday from the saphenous or tail vein using an EDTA-coated collection tube every 4 wk. Glucose levels were measured immediately after the puncture with an Accu-Check Aviva Nano glucose meter (Roche).

### Glucose and insulin tolerance tests

Mice were fasted for 6 h and blood glucose levels were measured from the tail vein before intraperitoneal injection. For glucose tolerance tests, MacHDAC7 and MacBlue mice were injected with 2 g/kg of D-glucose (AJAX), whereas *Hdac7*^-/-^ and *Hdac7*^+/+^ mice were injected with 1g/kg of glucose. For insulin tolerance tests, *Hdac7*^-/-^ and *Hdac7*^+/+^ mice were fasted for 6 h before being injected with 0.75 U/kg insulin (NovoRapid). Blood glucose levels were measured from the tail vein at the indicated time points with an Accu-Check Aviva Nano glucose meter (Roche). For the tests on MacHDAC7 and MacBlue mice, data were combined from two independent experiments.

### Histological assessment

Formalin-fixed mouse livers were processed for paraffin-embedded histology for Hematoxylin and Eosin (H&E) and Sirius Red staining. Unstained slides were processed for H&E staining using a DAKO autostainer. Sirius Red slides were processed using the Llewellyn modification of the Sirius Red technique^67^. In short, slides were stained with haematoxylin and then treated with Sirius Red solution for 1 h before washing in tap water for 10 min. Slides were imaged using a clinical microscope (BX43 or BX51, Olympus). Livers were scored using the NASH Clinical research Network (NASH-CRN) scoring system^68^. In this method, scores for steatosis, ballooning and lobular inflammation are summed to generate an overall score for likelihood of steatohepatitis.

### Immune profiling

MacBlue and MacHDAC7 mice, either at end-point of the chow diet versus HFHCHS diet trial (24 weeks: 30-34 weeks of age) or at 6 to 10 weeks of age, were anesthetized with isofluorane and 0.5-1 mL of blood was collected into heparinised tubes by cardiac puncture before cervical dislocation. Spleens were collected and processed as previously described^69^. In brief, spleens were weighed and then dissociated in 3 mL PBS 2% NCS 2 mM EDTA using a GentleMACS Dissociator tissue homogenizer with matching C tubes (Miltenyi Biotec, Macquarie Park, Australia) on “spleen 3” setting, twice. Spleen cells were counted either diluted 1:2 on a Coulter Counter (Beckman Coulter) or neat on a Mindray BC-5000 Vet Auto hematology analyser (Biomedical Electronics Co. LTD., China). Blood was counted neat on a Mindray BC-5000 Vet Auto hematology analyser. For immunophenotyping, spleen samples were stained in 50% CD16/CD32 hybridoma 2.4G2 supernatant for blocking and a fluorescent antibody cocktail containing anti-mouse CD45-Pacific Blue (BioLegend, Cat#103126, clone 30-F11, 1/200 dilution), anti-mouse CD3ε-FITC (BioLegend, cat# 100306, clone 145-2C11, 1/200 dilution), anti-mouse/human B220-APC-Cy7 (BioLegend, cat# 03224, clone RA3-6B2, 1/200 dilution) and anti-mouse/human CD11b-PerCP-Cy5.5 (BioLegend, cat# 101228, clone M1/70, 1/200 dilution) was used to measure numbers of T, B and total myeloid cells. 7-amino-actinomycin D (7AAD; ThermoFisher Scientific, cat# A1310, 1/10 dilution) or fixable viability stain (FVS)700 (BD Bioscience, cat# 564997, 1/10,000 dilution) was added to all stained samples for dead cell exclusion. Samples were analyzed on a Cytoflex (Beckman Coulter) flow cytometer equipped with 640 nm, 561 nm, 488 nm and 405 nm lasers. Uncompensated FCS files were analyzed using FlowJo 10.10 software, following compensation with single color antibody stains (Tree Star, Ashland, OR) and following post-hoc compensation with single color controls. The gating strategy used for flow cytometry analysis is shown in **Figure S5**.

### Glycogen quantification

The method used was previously described^70, 71^. Snap-frozen mouse livers were homogenised in 1 mL isolation buffer (50 mM Tris, pH 8, 150 mM NaCl, 2 mM EDTA, 50 mM NaF, 5 mM sodium pyrophosphate, Sigma-Aldrich). 20 μL homogenate was then digested in 22.5 μL acetate mix (1:8 mixture of 10% acetic acid (Sigma-Aldrich):200 mM sodium acetate buffer (Sigma-Aldrich) (pH4.5)) with 5 μL and amyloglucosidase ∼3260 U/mL, Megazyme) at 50 °C for 30 min. The same amount of undigested liver homogenates boiled for 10 min was used for correcting background glucose levels. Liver homogenates were then centrifuged at 20,000 *g* for 10 min and transferred to a 96-well plate. Serially diluted D-glucose was used for generating the calibration curve. 30 μL samples were then mixed with a reaction mix containing 150 μL of 200 mM tricine/KOH (pH 8) (Sigma-Aldrich) and 10 mM MgCl2 (Sigma-Aldrich), 1 μL of 112.5 mM NADP (NADP), 1 μL of 180 mM of ATP (NADP) and 0.5 μL glucose-6-phosphate dehydrogenase (Roche) to a volume of 200 μL. Absorbance at 340 nm was recorded for 20 min before as well as 30 min after the addition of hexokinase (Roche). Glycogen content was corrected for the weight of homogenised liver tissues.

### Insulin measurements

Blood collected from saphenous vein or by cardiac puncture was centrifuged at 10,000 rpm for 10 min. Concentrations of insulin in mouse serum or plasma were measured using a mouse insulin ELISA kit (Crystal Chem Mouse Insulin ELISA Kit, 90080), following the manufacturer’s instruction. Standard solutions and samples were added in duplicate into the wells included in the kits and incubated for 2 h at 4 °C. Each well was then washed thoroughly.

Next, enzyme conjugate solution was added and incubated for 30 min at room temperature, followed by another washing step. Enzyme substrate solution was immediately added and incubated for 40 mi at room temperature in the dark. Following that, enzyme stop solution was added to each well to stop the reaction. Finally, the absorbance at A450 and A630 was measured using a microplate reader. Sample insulin concentrations were calculated based on a standard curve generated using known insulin concentrations from the standard solutions included in the kits.

### Cell culture

Bone marrow cells were collected from femurs and tibias of mice and cultured in RPMI media supplemented with 10% FBS, 2mM L-glutamine, 50 U/mL penicillin and 50 μg/mL streptomycin (Gibco) in the presence of 150 ng/mL recombinant human macrophage colony-stimulating factor-1 (Protein Expression Facility, UQ) for 6 days. Differentiated BMMs were harvested in DPBS (Thermo Fisher Scientific) and resuspended in culture media for stimulation with indicated concentrations of LPS (*Salmonella enterica* serotype Minnesota, L2137, Sigma-Aldrich). To induce IL-1β processing and secretion, 5 μg/mL nigericin (N7143, Sigma-Aldrich) was added to cells 1 h prior to harvesting culture supernatants. Cell culture supernatants were then collected, centrifuged at 1,000 *g* for 5 min and stored at -80 °C until analysis.

### Cytokine profiling

Cell culture supernatants and mouse plasma were diluted in culture media and DPBS, respectively. CCL2 and IL-1β concentrations were measured using MCP-1/CCL2 Mouse Uncoated ELISA and IL-1beta Mouse Uncoated ELISA Kits (mouse CCL2: 88-7391-77, mouse IL-1β: 88-7013-77, Invitrogen) or with a Mouse CCL2/JE/MCP-1 DuoSet ELISA kit (DY479, R&D SYSTEMS), as per the manufacturer’s manuals.

### Quantification of gene expression

To extract RNA from mouse livers, fresh tissues were frozen in lysis buffer from an ISOLATE II RNA Mini kit (BioLine) at -80 °C. Snap-frozen tissues from naïve mice were homogenised in lysis buffer. RNA purification was performed following the manufacturer’s instructions. An off-column DNA digestion was performed using DNAse I (Invitrogen), as per manufacturer’s instructions. RNA was reverse transcribed to cDNA using oligo dT and SuperScript III Reverse Transcriptase (Invitrogen). Quantitative PCR (qPCR) was performed in 384-well plates using cDNAs, SYBR Green PCR Master Mix (Applied Biosystem), and a ViiATM 7 Real-Time PCR System (Applied Biosystems). mRNA levels of individual genes, relative to the housekeeping gene hypoxanthine phosphoribosyl transferase (*Hprt*), were calculated using the ΔCt method^72^. Sequences of primers used for qPCR are listed in **Table S2**.

### Western blotting

Differentiated BMMs were washed once with DPBS and lysed in RIPA buffer (50 mM Tris-HCl, 150 mM NaCl, 0.1% SDS, 1% Sodium deoxycholate, 1% NP-40) supplemented with protease inhibitor cocktail (cOmpleteTM, Roche) and phosphatase inhibitors (PhosStopTM, Roche). Cell lysates were homogenised by passing through a 27G needle. Equal amounts of homogenates were boiled and subjected to electrophoresis using pre-cast 4-12% Bis-Tris NuPAGE gels in NuPAGE MOPS SDS running buffer (Invitrogen). After separation, proteins were transferred to a nitrocellulose membrane and immunoblotted for endogenous HDAC7 using an anti-HDAC7 antibody (Cell Signaling Technology, 1/1000) followed by an HRP-conjugated goat anti-rabbit IgG (Cell Signaling Technology, 0.026 µg/mL) and Tubulin as a loading control using a fluorescent Rhodamine anti-Tubulin antibody (Bio-Rad, 1/5000).

### Quantification of hepatic *HDAC7* mRNA levels in human CLD

Hepatic *HDAC7* mRNA levels were quantified in a cohort of 69 people with CLD of mixed aetiologies. Diagnosis, disease scoring and sample stratification has previously been described^51^. *HDAC7* mRNA levels in people with early (stages 0-2) and advanced (stages 3-4) CLD were quantified by RNA sequencing, as previously described^51^. In brief, sequencing reads were mapped to the human hg19 genome using STAR aligner^73^ and CPM values of transcript reads were obtained using the edgeR package^74^. Genes were filtered with a CPM threshold of over 10 in at least 20 samples and subjected to normalisation using the Trimmed Mean of M-values method using edgeR^75^. Differential gene expression was then analysed by likelihood-ratio test, with multiple testing performed using Benjamini-Hochberg corrections^76^. FDR values of less than 0.05 were considered significantly different.

### Spatial transcriptomics analyses

Liver tissues were harvested from two male MacBlue and two male MacHDAC7 mice and immediately fixed in 4% paraformaldehyde for 24 h at 4°C. Fixed tissues were processed, embedded in paraffin, and sectioned at 5 µm thickness. Sections were mounted onto Xenium V1 slides (10x Genomics) and processed according to the manufacturer’s protocol for Xenium Prime In Situ Gene Expression with Cell Segmentation Staining (10x Genomics, CG000760 RevC). Briefly, slides underwent deparaffinization and decrosslinking, followed by priming hybridization, RNase treatment and polishing. The Xenium 5K Ms PTP Panel Probes (10x Genomics, Cat#2001227), as well as an add-on custom probe panel targeting 50 genes, including key macrophage, inflammatory, metabolic, and fibrosis-associated transcripts (**Table S3**), was hybridised to the tissue sections overnight. Following probe ligation, rolling circle amplification and cell segmentation staining steps, slides were loaded onto Xenium Analyzer for signal detection and image acquisition. Raw image data were processed using the Xenium Onboard Analysis pipeline, and decoded transcripts were mapped back to the tissue at the single cell level. Data were analysed using Seurat workflow (Seurat version 5.1.0), including quality control filtering, and gene expression quantification in R (version 4.4.0). EdgeR (version 4.4.0) was used to perform differential gene expression analysis on MacBlue versus MacHDAC7 hepatic cells that were positive for both *eCFP* and *Hdac7* mRNAs and the results were visualised on a heatmap. Genes overlapping with previously published CLD-associated signatures were further examined for spatial co-localisation with bivariate Moran’s Index (from esda package version 2.2.1) using custom python scripts and were visualised with spatial feature plots.

### Statistical analyses

Data from *in vivo* experiments are combined from one or two independent trials with each individual mouse representing one independent data point. For cell culture experiments, means of experimental replicates were determined, with data from independent mice or experiments being combined for statistical analysis. Statistical analyses were performed using Prism 9 or 10 (GraphPad). Comparisons between two groups were performed using a two-tailed unpaired Student’s *t*-test or Mann Whitney U test as indicated. For datasets involving at least three variables, linear regression, ordinary or repeated measures two-way ANOVA tests were performed followed by multiple comparisons, as indicated in individual figure legends. Differences with confidence value of 95% and above (*p* < 0.05) were considered statistically significant.

## Acknowledgements

This work was supported by a National Health and Medical Research Council of Australia (NHMRC) Ideas grant (1184885 to MJS and DR), an NHMRC Investigator grant to MJS (1194406), and an Australian Infectious Disease Research Centre seed grant to CRE, MJS, JAE and DR. DK was supported by an Institute for Molecular Bioscience (IMB) Fellowship and an NHMRC Ideas grant (2012772). DPF was supported by an NHMRC Investigator grant (2009551) and an ARC Centre of Excellence grant (CE200100012). SSB was supported by a Swiss National Science Foundation Postdoc Mobility Fellowship (P2BEP3_191800) and a Novartis Foundation for Medical-Biological Research Fellowship (21C133). KMI was supported by an NHMRC Ideas grant (2012793).

## Author contributions

Conceptualisation, Y.W., M.A.S., M.J.S., D.K.; Methodology, G.C.M., S.K., K.S.G., Z.L., S.S.B., K.M.I; Investigation, Y.W., D.R., K.D.G., P.P., K.B., G.C.M., Z.X., Yu.W., E.N.T., J.E.B.C., R.A., K.S.G., J.D.A., D.K.; Resources, G.C.M., J.A.E., C.R.E, A.D.C., E.E.P., J-P.L., Q.N.; Writing- original draft, Y.W., M.J.S., D.K.; Writing- Review/Writing, Y.W., G.C.M., P.P., S.S.B., D.P.F., K.M.I., M.A.S., M.J.S., D.K.; Supervision, K.S., D.P.F., A.D.C., K.M.I., M.A.S., J-P.L., Q.N., M.J.S., D.K.; Funding acquisition, D.R., C.R.E, J.A.E., M.J.S.

---

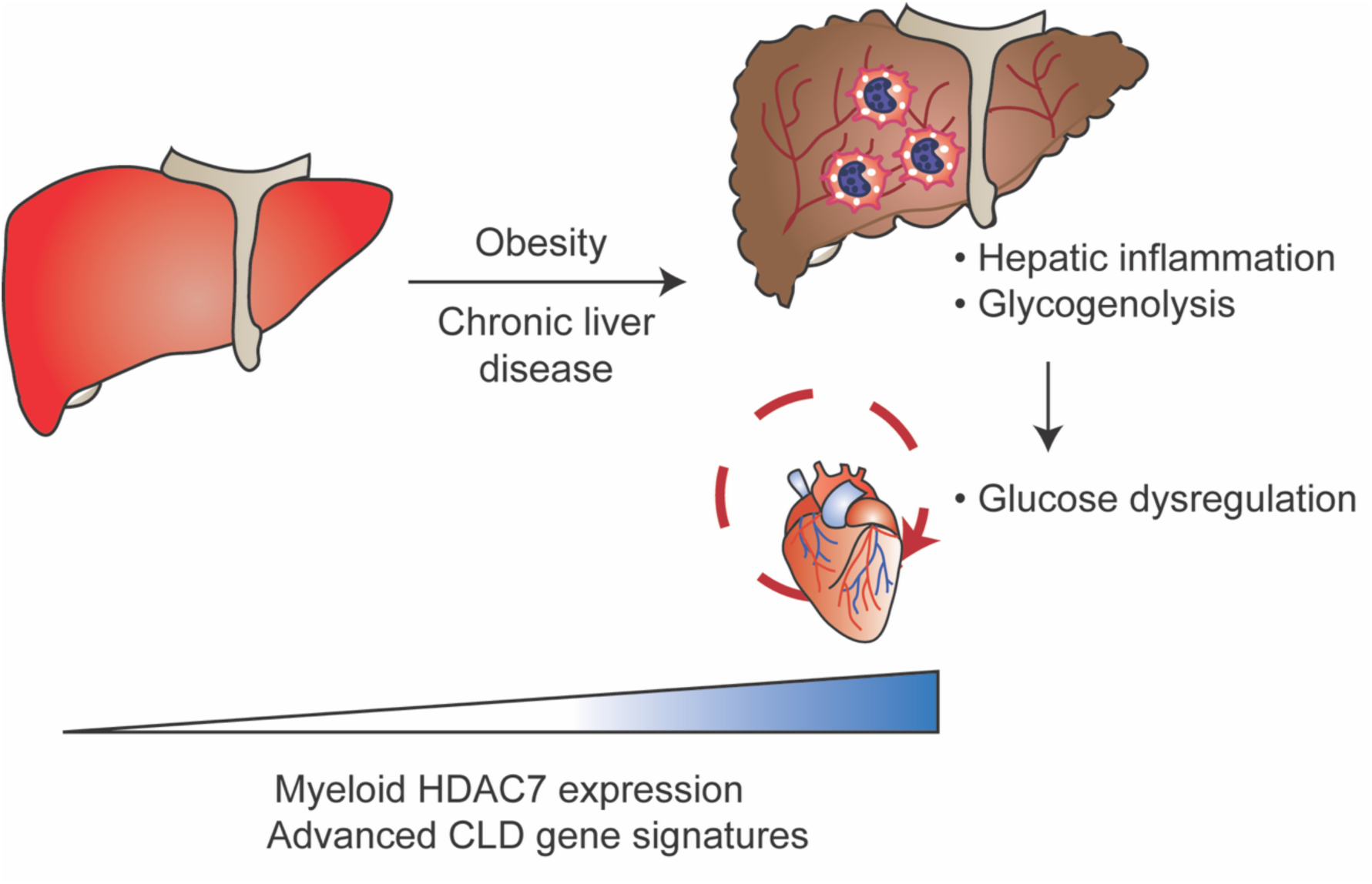

This study explored the role of the class IIa histone deacetylase HDAC7 in hepatic inflammation and metabolic disease. Genetic approaches in mice revealed that myeloid HDAC7 drives hepatic inflammation and gene signatures associated with advanced chronic liver disease. Myeloid HDAC7 also dysregulates systemic glucose metabolism during diet-induced obesity in mice.

