## Supplementary Material for "Myeloid HDAC7 drives liver inflammation and systemic glucose dysregulation during diet-induced obesity"

### SUPPLEMENTARY INFORMATION

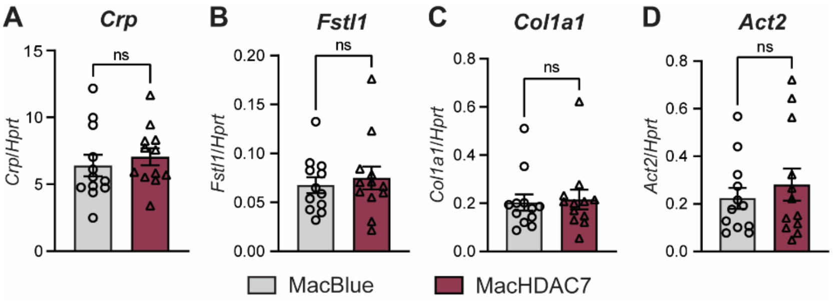


**Supplementary Figure S1. Hepatic mRNA expression of inflammatory and fibrotic genes in MacHDAC7 mice.**

Hepatic mRNA levels of inflammation marker *Crp* (**A**), and fibrotic markers *Fstl1* (**B**), *Col1a1* (**C**) and *Act2* (**D**) in livers from 9-22 wk old MacHDAC7 and control MacBlue mice fed a chow diet (same cDNA samples as used in Figure 1). Data, expressed relative to levels of *Hprt* mRNA, are mean ± SEM from n=12 mice (6 male and 6 female) for each group. Statistical analyses were performed using Student's *t*-test or (ns, not significant).

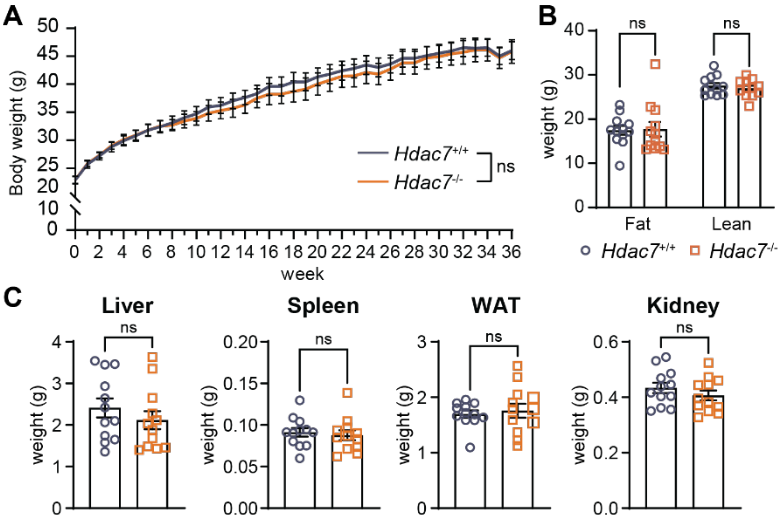


**Supplementary Figure S2. Effect of myeloid *Hdac7* deletion on body weight in mice fed a chow versus HFHCHS diet.**

8-10 wk old male *Hdac7*$^{fl/fl}$ (*Hdac7*$^{+/+}$) and *Hdac7*$^{fl/fl}$ *LysM*$^{Cre}$ (*Hdac7*$^{-/-}$) mice were fed a HFHCHS diet for 36 wk. **A.** Body weight was measured weekly. **B.** Fat and lean mass measured by EchoMRI at wk 35. **C.** Total organ weights for liver, spleen, epididymal white adipose tissue (WAT) and kidney were measured at the termination of the study. Data shown are mean ± SEM of 12 mice from each group. Statistical analysis was performed using linear regression (**A**), repeated measures two-way ANOVA followed by Sidak's multiple comparison (**B**) or Student's *t*-test (**C**) (ns, not significant).

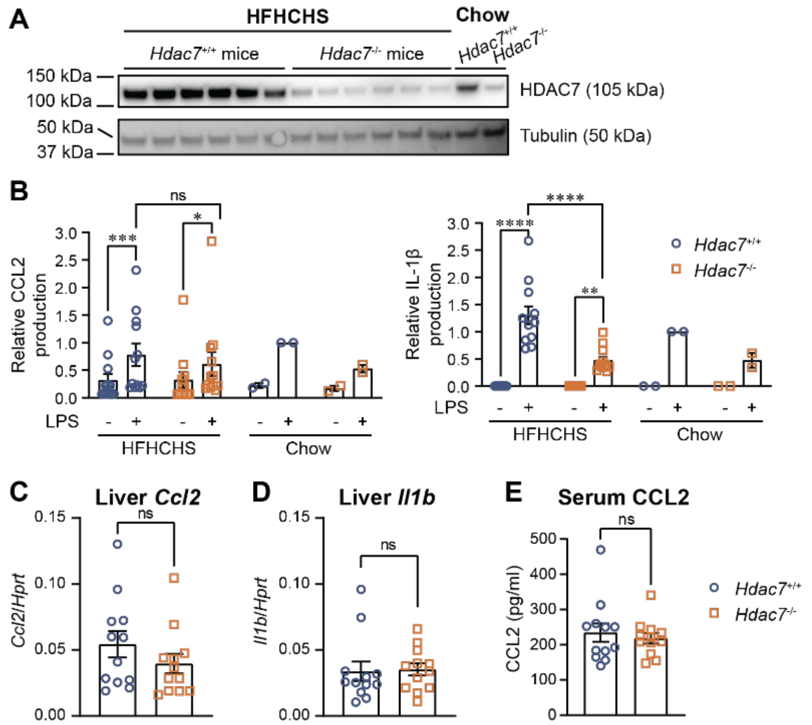


**Supplementary Figure S3. Myeloid *Hdac7* deletion reduces inflammatory responses in macrophages but does not affect hepatic inflammatory gene expression or serum CCL2 in mice fed a HFHCHS diet.**

**A.** Bone marrow cells from *Hdac7*$^{+/+}$ and *Hdac7*$^{-/-}$ mice on the HFHCHS diet were collected and differentiated in the presence of CSF-1 for 7 days and lysed, after which protein levels of HDAC7 were determined by immunoblotting, with Tubulin as a loading control (n=6 per group). BMMs from a pair of 12 wk old *Hdac7*$^{+/+}$ and *Hdac7*$^{-/-}$ mice on a chow diet were used as a positive control. **B.** Differentiated BMMs from mice of indicated genotypes on a HFHCHS diet were stimulated with LPS (0.5 ng/ml) for 4 h. CCL2 (left) and nigericin-induced IL-1β secretion (right) in culture supernatants were measured by ELISA (left hand side of graphs). BMMs from 12 wk old mice on a chow diet were used as inter-assay control (right hand side of graphs). Relative CCL2 and IL-1β production was normalised to the LPS-inducible production from BMMs of control *Hdac7*$^{+/+}$ mice on the chow diet. Data shown are mean ± SEM of 12 mice for each group and were analysed using repeated measures two-way ANOVA followed by Bonferroni's correction (* $p < 0.05$; *** $p < 0.001$; **** $p < 0.0001$). **C-E.** Quantification of inflammatory mediators in the liver and circulation. mRNA levels of *Ccl2* (**C**) and *Il1b* (**D**) (relative to *Hprt*) were measured from freshly homogenised frozen livers. CCL2 levels in mouse sera was measured by ELISA (**E**). Data shown are mean ± SEM from 12 mice for each group and were analysed using Student's *t*-test (ns, not significant).

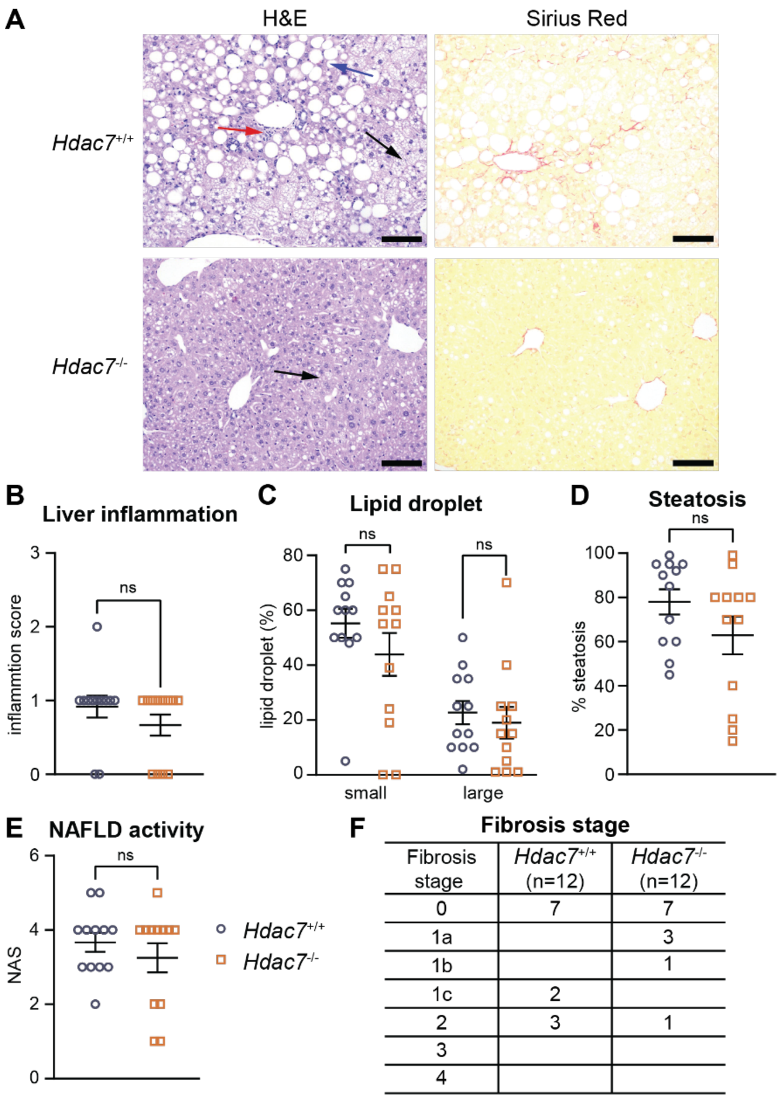


| Fibrosis stage | *Hdac7*$^{+/+}$ (n=12) | *Hdac7*$^{-/-}$ (n=12) |
|---|---|---|
| 0 | 7 | 7 |
| 1a | | 3 |
| 1b | | 1 |
| 1c | 2 | |
| 2 | 3 | 1 |
| 3 | | |
| 4 | | |

**Supplementary Figure S4. Histological assessment of livers from *Hdac7*$^{+/+}$ and *Hdac7*$^{-/-}$ mice.** At termination of the study, mice were fasted for 12 h and euthanised. **A.** Liver pathology was assessed after H&E (left) and Siriud Red staining (right). Arrows indicate microvesicular steatosis (black), macrovesicular steatosis (blue), and a focus of inflammation within the lobule (red), scale bar: 100 µm. **B-F.** Quantification of lobular inflammation (**B**), lipid droplet counts (**C**), steatosis (**D**), NAFLD activity score (NAS, **E**) and fibrosis stage (**F**) as per the NASH clinical research network scoring system after H&E staining. Data shown represent mean ± SEM of 12 mice per group and were analysed using Student's *t*-test (**B**, **D-E**) or two-way ANOVA followed by Sidak's multiple comparison **(C)** (ns, not significant).

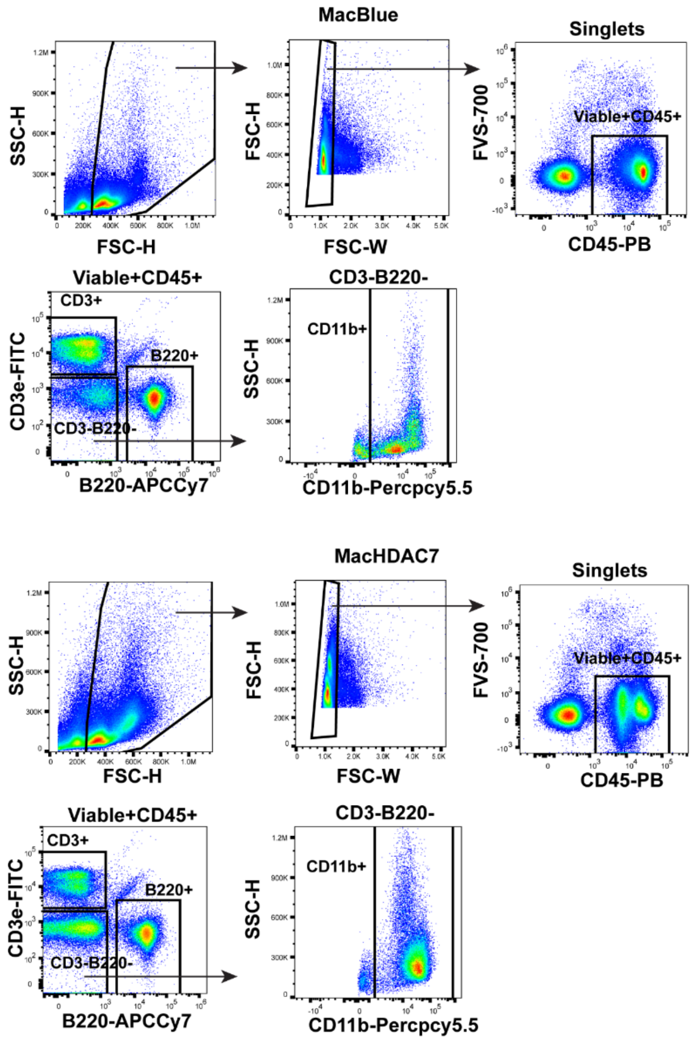


**Supplementary Figure S5. Flow cytometry gating strategy to identify CD3$^+$ T cells, B220$^+$ B cells and CD11b$^+$ myeloid cells.**

This example is taken from a spleen sample from a MacBlue (top) and a MacHDAC7 mouse (bottom). Singlet cells were gated according to forward and side scatter and then gated for viable (FVS-negative) CD45$^+$ leukocytes. CD3$^+$ T and B220$^+$ B cells were gated on viable+CD45+ leukocytes. CD3/B220$^-$ cells were gated to identify CD11b$^+$ myeloid cells.

**Supplementary Table S1. Fibrosis stage of livers from MacBlue and MacHDAC7 mice.**

| Fibrosis stage | Chow | | HFHCHS | |
|---|---|---|---|---|
| | MacBlue (n=6) | MacHDAC7 (n=6) | MacBlue (n=14) | MacHDAC7 (n=14) |
| 0 | 6 | 6 | 5 | 9 |
| 1a | | | 3 | 3 |
| 1b | | | 1 | 1 |
| 1c | | | | |
| 2 | | | 2 | |
| 3 | | | 3 | 1 |
| 4 | | | | |

After H&E staining, fibrosis stage of livers from MacBlue and MacHDAC7 mice fed on the HFHCHS diet (from Figure 3) was scored according to the standards of NASH clinical research network scoring system. Numbers of mice at each stages of fibrosis is indicated in individual table cells.

**Supplementary Table S2. Primer sequences used in this study.**

| Gene | Forward (5'-3') | Reverse (5'-3') |
|---|---|---|
| *Hdac7* | CGCAGCCAGTGTGAGTGTCT | AGTGGGTTCGTGCCGTAGAG |
| *Ccl2* | CCCACTCACCTGCTGCTACTCA | GCTTCTTTGGGACACCTGCTG |
| *Il1b* | GAAGTTGACGGACCCCAAAA | GCCTGCCTGAAGCTCTTGTT |
| *Mmp9* | TTGAGTCCGGCAGACAATCC | CCTTATCCACGCGAATGACG |
| *Crp* | GGCCAGATGCAAGCATCATC | CTGGAGATAGCACAAAGTCCCAC |
| *Fstl1* | CAACCCATCCTTCAACCCTCCT | ATTCTTTCCATCACAGGTCAT |
| *Col1a1* | CGTATCACCAAACTCAGAAG | GAAGCAAAGTTTCCTCCAAG |
| *Act2* | CATCTTTCATTGGGATGGA | TTAGCATAGAGATCCTTCCTG |
| *Hprt* | GTTGGATACAGGCCAGACTTTGTTG | GAGGGTAGGCTGGCCTATAGGCT |
| *Adgre1* | CTCTGTGGTCCCACCTTCAT | GATGGCCAAGGATCTGAAAA |
| *Clec4f* | TTTTGTGGTGGCTTCACAGC | TCGTGTCCAGAGTTGTTGCT |

**Supplementary Table S3. Custom probe panel for spatial transcriptomic analysis of mouse livers**

| | | | | |
|---|---|---|---|---|
| *Acta2* | *Dgat2* | *Hdac11* | *Aff3* | *Adk* |
| *Clec4f* | *Pgd* | *Tmem68* | *S100a4* | *Dapl1* |
| *Vsig4* | *Plin3* | *Nkg7* | *Id3* | *Fgl2* |
| *Crp* | *Ehf* | *Ccl5* | *Ccl4* | *Sesn3* |
| *Il1b* | *Slc30a4* | *Klf2* | *Rgs10* | *Mt1* |
| *Il6* | *Ptges2* | *Klf6* | *Asb2* | *ECFP* |
| *Spp1* | *Plin4* | *Cd7* | *Cebpb* | *LdLV9.34.2.208810.1* |
| *Hdac7* | *Hilpda* | *Zfp683* | *S100a6* | *V5RNA* |
| *Slc30a1* | *G0s2* | *Slamf7* | *Lgals1* | *CD45.2_WT_93* |
| *Slc39a14* | *Hdac10* | *Nsg2* | *Lgals3* | *CD45.2_ALT_93:TC* |
